# Sox2-RNA mechanisms of chromosome topological control in developing forebrain

**DOI:** 10.1101/2020.09.22.307215

**Authors:** Ivelisse Cajigas, Abhijit Chakraborty, Madison Lynam, Kelsey R Swyter, Monique Bastidas, Linden Collens, Hao Luo, Ferhat Ay, Jhumku D. Kohtz

**Affiliations:** Department of Pediatrics, Northwestern University, Feinberg School of Medicine, Department of Human Molecular Genetics, Stanley Manne Children’s Research Institute 2430 N Halsted, Chicago, IL 60614; Centers for Autoimmunity and Cancer Immunotherapy, La Jolla Institute for Immunology, 9420 Athena Circle, La Jolla, CA, 92037, USA; School of Medicine, University of California San Diego, 9500 Gilman Drive, La Jolla, CA 92093, USA

**Keywords:** lncRNA, chromosome 3D structure, architectural proteins, enhancers, epigenetics, forebrain development, ribonucleoprotein complex

## Abstract

Precise regulation of gene expression networks requires the selective targeting of DNA enhancers. The *Evf2* long non-coding RNA regulates *Dlx5/6* ultraconserved enhancer*(UCE)* interactions with long-range target genes, controlling gene expression over a 27Mb region in mouse developing forebrain. Here, we show that *Evf2* long range gene repression occurs through multi-step mechanisms involving the transcription factor Sox2, a component of the *Evf2* ribonucleoprotein complex (RNP). *Evf2* directly interacts with Sox2, antagonizing Sox2-dependent *Dlx5/6UCE* activation*. Evf2* regulates Sox2 binding at key sites, including the *Dlx5/6eii* shadow enhancer and *Dlx5/6UCE* interaction sites. *Evf2* differentially targets RNP-associated Sox2 protein pools (PPs), redirecting Sox2-PPs to one repressed gene at the expense of the other. Co-regulation of *Dlx5/6UCE*intrachromosomal interactions by *Evf2* and Sox2 reveals a role for Sox2 in chromosome topology. We propose that RNA organizes RNPs in a subnuclear domain, regulating both long-range *UCE* targeting and activity through Sox2-RNP sequestration and recruitment.

## Introduction

Ultraconserved elements (UCEs) were identified as 200 base pair (or greater) segments of 100% DNA conservation between humans, mice and rats, many associated with key developmental regulators (Bejerano et al., 2004; Sandelin et al., 2004; Woolfe et al., 2005). Removal of UCEs in mice initially suggested that UCEs are dispensable (Ahituv et al., 2007). However, removal of UCE sequences near developmental regulators Arx, Gli, and Shox2 cause neurological and growth defects (Dickel et al., 2018; Osterwalder et al., 2018), and limb defects (Nolte et al., 2014), revealing specific developmental roles. Transcription of UCE sequences and enhancer-regulating activity of UCE transcripts (Calin et al., 2007; Feng et al., 2006), together with the identification of genome-wide scale enhancer transcripts with enhancer-like activities (Orom et al., 2010; Orom and Shiekhattar, 2011), support mechanistic and functional diversity of RNA regulatory roles (Rinn and Chang, 2020).

Our studies on the *Evf2* ultraconserved enhancer lncRNA (*Dlx5/6UCE*-lncRNA, overlapping with *Dlx6OS1*) support complex RNA regulatory roles for UCE sequences during embryonic forebrain development (Berghoff et al., 2013; Bond et al., 2009; Cajigas et al., 2018; Cajigas et al., 2015; Feng et al., 2006). *Evf2* is a 3.7kb, spliced and polyadenylated lncRNA, containing *Dlx5/6UCE* sequences responsible for enhancer-regulating activities, in *trans* (Feng et al., 2006). *Evf2* controls an embryonic brain interneuron gene regulatory network (GRN), adult hippocampal and cortical circuitry defects, and seizure susceptibility in adult mice (Bond et al., 2009; Cajigas et al., 2018; Feng et al., 2006). *Evf2* positively and negatively regulates gene expression through *cis* (same chromosome as *Evf2* expression site) and *trans* (different chromosome) mechanisms (Berghoff et al., 2013; Cajigas et al., 2018). Mechanisms of *Evf2* gene activation and repression are distinguished by different requirements of the RNA, with the *5’*-UCE containing region of the lncRNA controlling gene repression and the 3’ end controlling gene activation (Cajigas et al., 2018).

*Evf2* RNA cloud-formation is similar to lncRNAs that regulate dosage compensation (*Xist* (Brockdorff et al., 1992; Brown et al., 1992)) and imprinting (*Kcnq1ot1* (Pandey et al., 2008; Redrup et al., 2009)). *Evf2* assembles a ribonucleoprotein complex (RNP^87^) containing 87 functionally diverse proteins, including transcription factors (TFs: Dlx1, Sox2), chromatin remodelers (Smarca4, Smarcc2, Smarcb1), regulators of chromosome topology (Smc1, Smc3), and lamin B1 (Cajigas et al., 2015) (Fig 1A). *Evf2*-dependent gene regulation across a 27MB region of mouse chromosome 6 (*chr6*) is characterized by recruitment of individual RNP proteins (TFs, chromatin remodelers, cohesin), and regulation of epigenetic modifications at key DNA regulatory sites, including the *Dlx5/6UCE*.

**Figure 1.**
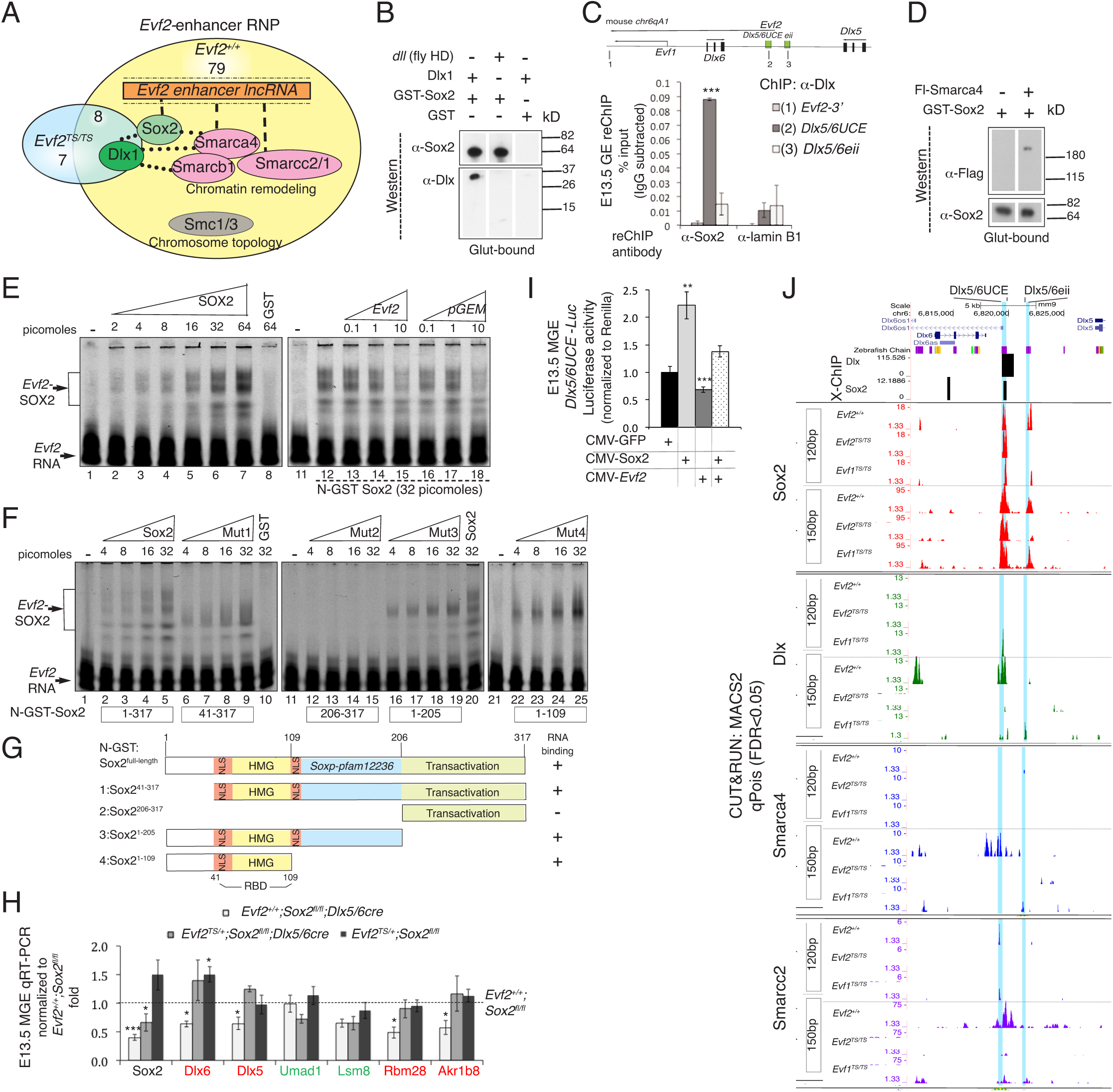
*Evf2* repression through Sox2 antagonism in mouse developing forebrain. **A.** The *Evf2* UCE lncRNA and Sox2 protein are RNA/protein scaffolds for multiple RNPs, including chromatin remodelers (SmarcA4/C2/C1) and cohesin (Smc1/3) in E13.5 mouse ganglionic eminence (GE). Sox2 is one of 79 proteins complexed with Dlx1 in the presence of *Evf2* (*Evf2^+/+^*), and not detected in GE lacking *Evf2* (*Evf2^TS/TS^*). **B.** Western analysis shows direct binding of GST-Sox2 to mouse Dlx1, but not GST or Dll (fly homeodomain fragment). **C.** Dlx-Sox2 complexes are detected at the *Dlx5/6UCE* by ChIP-reChIP (first anti-Dlx, second anti-Sox2) from GE crosslinked chromatin. **D.** Western analysis shows that GST-Sox2 directly binds flag-tagged Smarca4. **E-G.** RNA electrophoretic mobility shift assays (REMSAs) using infrared labelled *Evf2* RNA as a probe. **E**. Sox2 binding to *Evf2* RNA at picomolar concentrations is promiscuous, as indicated by pGEM RNA competition with *Evf2* RNA for binding to Sox2. **F.** The Sox2 RNA binding domain is narrowed to the Sox2 high mobility group domain (HMG) and adjacent nuclear localization (NLS) regions. **G**. Schematic of Sox2 mutants tested in REMSAs. **H.** TAQman qRT-PCR analysis of E13.5 medial ganglionic eminences (MGE) from mice lacking Sox2 in interneuron progenitors (*Evf2^+/+^*;*Sox2^fl/fl^;Dlx5/6cre*), additional loss of one copy of (*Evf2^TS/+^*;*Sox2^fl/fl^;Dlx5/6cre*), loss of one copy of *Evf2* (*Evf2^TS/+^*;*Sox2^fl/fl^*), normalized to wildtype (*Evf2^+/+^*;*Sox2^fl/fl^*). Sox2 loss reduces *Evf2*-repressed target genes in red (*Dlx5/6*, *Rbm28* and *Akr1b8*) white bars, rescued by loss of one copy of *Evf2* (grey bars). n=3-6/genotype, **I.** Luciferase assays of E13.5 MGE using a *Dlx5/6UCE*-luciferase reporter (*Dlx5/6UCE*-Luc) shows that Sox2 activation of *Dlx5/6UCE* is inhibited by *Evf2* repressor activity. n=6/condition, replicated. **J.** CUT&RUN native ChIPseq binding profiles of *Evf2*-RNPs (Sox2, Dlx, Smarca4, Smarcc2) in *Evf2^+/+^*, *Evf2^TS/TS^*, *Evf1^TS/TS^* GE. *Dlx5/6* intergenic enhancers (*Dlx5/6UCE* and *Dlx5/6eii* shadow enhancer) are highlighted in blue. 120bp and 150 bp profiles show MACS2 validated peaks (FDR <0.05), n=2-4/genotype. Comparisons between *Evf2* loss (*Evf2^TS/TS^*) and truncation (*Evf1^TS/TS^*) mutants identify *Evf2* 5’ vs. 3’ Sox2 and Smarcc2 differentially regulated sites. In G-I: Student’s *t*-test, *p<0.05, **p<0.01***p<0.001, error bars indicate SEM (% standard error of the mean).

The identification of the TF Sox2, a component of *Evf2*-RNP (Cajigas et al., 2015), raised questions about its role in lncRNA-mediated gene regulation. Sox2 is a well-characterized pioneer transcription factor (Dodonova et al., 2020) that maintains pluripotency through lineage-specific gene repression (Avilion et al., 2003; Takahashi and Yamanaka, 2006). Sox2 associates with lncRNAs involved in pluripotency and neuronal differentiation (Guo et al., 2018; Ng et al., 2013; Ng et al., 2012), and binds both DNA and RNA through its high mobility group domain (HMG) (Holmes et al., 2020). Crystal structures of HMG-POU-DNA ternary complexes support Sox2 multivalency and concentration-dependent enhancer regulation (Remenyi et al., 2003; Remenyi et al., 2004; Williams et al., 2004). In this report, we show that *Evf2* regulates *Dlx5/6UCE* targeting and activity through mechanisms involving the *Evf2*-RNP Sox2, revealing multi-step contributions of TF-RNA interactions. Sox2 and *Evf2*-RNP proteins NONO, Smarcc2, and Smc1/3 colocalize with *Evf2* RNA clouds, subnuclear domains that we have termed protein pools (PPs). *Evf2* controls PP targeting and sizes to repressed genes, and recruits individual *Evf2*-RNP proteins to key DNA regulatory sites, including *Dlx5/6* intergenic enhancers and enhancer-chromosome interaction sites. We propose that the *Evf2*-RNP functions as an organizer, distributing critical regulatory proteins to key DNA regulatory sites, that through indirect and direct interactions with RNP, control both *UCE* targeting and activity.

## Results

### *Evf2* gene repression through Sox2 antagonism

The *Evf2* RNA cloud is a scaffold for the assembly of the *Evf2*-RNP, directly binding chromatin remodelers Smarca4 and Smarcc2/1 through promiscuous RNA-protein interactions, and indirectly to the Dlx homeodomain TF. Previous work shows that Smarca4 bridges the *Evf2* RNA with the non-RNA binding protein Dlx1, and also binds RNA binding proteins within the RNP (Fig 1A, Cajigas et al. 2015). In order to investigate the role of individual *Evf2-RNP* ^87^ proteins in gene activation and repression, we addressed the role of the pioneer TF Sox2 in *Evf2* regulated gene expression. First, we studied Sox2 interactions with *Evf2*-RNP components. In the absence of *Evf2*, there is ∼25% decrease in total Sox2 protein levels (Fig S1A, B), and ∼50% increase in Sox2 RNA (Fig S1C), supporting the involvement of both transcriptional and post-transcriptional control mechanisms. Consistent with Sox2 directly binding Dlx1 (Fig 1B), ChIP-reChIP shows that Sox2 and Dlx simultaneously bind *Dlx5/6UCE* (Fig 1C). In addition, Sox2 directly binds Smarca4 (Fig 1D), supporting multiple protein partners within the *Evf2*-RNP. RNA electrophoretic mobility shift assays (REMSAs) show that *Evf2* RNA binding to Sox2 is promiscuous, requiring Sox2 amino acids 41-109 (high mobility group [HMG] DNA binding domain and N-terminal nuclear localization signal [NLS], Fig 1E-G). Thus, Sox2 is similar to Smarca4, potentially functioning as a protein bridge between non-RNA binding proteins in the *Evf2-RNP* and *Evf2* RNA (Fig 1A). Sox2 RNA binding has been recently characterized, showing the requirement for the HMG domain for promiscuous Sox2-*ES2* lncRNA interactions (Holmes et al., 2020)

In order to determine whether Sox2 contributes to *Evf2*-dependent gene activation or repression, we analyzed gene expression in E13.5GE (*Sox2^fl/fl;^Dlx5/6cre+*), a genetic model in which floxed *Sox2* (Shaham et al., 2009) allows removal of Sox2 from *Dlx5/6+* GABAergic progenitors (Monory et al., 2006). These comparisons show that Sox2 activates *Evf2*-repressed target genes (in red), but does not affect *Evf2*-activated target genes (green) (Fig 1H). Loss of one copy of *Evf2* from *Sox2^fl/fl;^Dlx5/6cre+* E13.5GE (*Evf2^TS/+^;Sox2^fl/fl;^Dlx5/6cre*) rescues effects of Sox2 loss on repressed target genes (Fig 1H). Furthermore, *Dlx5/6UCE*-luciferase reporter assays show that *Evf2* antagonizes Sox2 activation of *Dlx5/6UCE* activity in E13.5GEs (Fig 1I), supporting a mechanism of *Evf2*-Sox2 antagonism during gene repression.

### *Evf2* regulates Sox2 binding to the *Dlx5/6eii* shadow enhancer, and Dlx, Smarca4 and Smarcc2 binding to *Dlx5/6UCE*

In order to determine if *Evf2/Sox2* regulation of *Dlx5/6UCE* activity involves Sox2 recruitment to the *Dlx5/6* intergenic enhancers, we used the native ChIPseq method CUT&RUN (Fig 1J red peaks indicate Sox2 binding). In the CUT&RUN method, sequencing of <120BP and >150bp fragments distinguish between proteins directly bound to DNA (less than 120bp), and indirect binding through protein-protein interactions (more than 150bp) (Meers et al., 2019a; Meers et al., 2019b). Analysis of >150bp CUT&RUN peaks has the potential to detect proteins associated with the large *Evf2-RNP*, not previously possible using crosslinked ChIPseq (X-ChIP) methods. CUT&RUN analysis (Methods) shows that while *Evf2* recruits *Evf2*-RNPs Dlx, Smarca4 and Smarcc2 to *Dlx5/6UCE,* Sox2 binding to Dlx5/6UCE is *Evf2*-independent (Fig 1J). Instead *Evf2* recruits Sox2 binding to *Dlx5/6eii*, a shadow enhancer (Furlong and Levine, 2018; Zerucha et al., 2000) located adjacent to *Dlx5/6UCE* and regulated by both Dlx and *Evf2* in *trans* assays, similarly to *Dlx5/6UCE* (Feng et al., 2006; Zerucha et al., 2000). The significance of *Evf2* mediated Sox2-*Dlx5/6eii* recruitment with respect to gene repression is supported by rescue in *Evf1^TS/TS^*, a genetic model where *Evf2* repression is also rescued (Cajigas et al., 2018) (Fig 1J). Analysis of 150bp profiles indicates that loss of *Evf2*-RNP (Dlx, Smarca4, Smarcc2) binding to *Dlx5/6UCE* in *Evf2^TS/TS^* (expressing only *Evf2*-3’) is not rescued in *Evf1^TS/TS^* (expressing only *Evf2*-5’). However, 120bp profiles support partial rescue of Smarcc2 direct binding to *Dlx5/6UCE*, and increased binding of Dlx in *Evf1^TS/TS^*. These data suggest *Evf2-5’* and -3’ regions are required for Dlx, Smarca4/Smarcc2 recruitment to *Dlx5/6UCE* through interactions with the larger *Evf2*-RNP complex, linking these events to gene activation. Furthermore, a role for *Evf2-5’* regulated Sox2 indirect and direct binding to the *Dlx5/6eii* shadow enhancer further supports Sox2 recruitment with gene repression, building on previous work showing that the *Evf2-5’* region is sufficient for repression (Cajigas et al., 2018). Together, these data support that *Evf2-5’* and 3’ differentially contribute to direct and indirect recruitment in a site-specific and RNP-dependent manner, linking individual recruitment events to gene repression and activation.

### Linking *Evf2* and Sox2 regulated *Dlx5/6*UCE targeting, and *Evf2* regulated RNP recruitment

Previous work showed that *Evf2-Dlx5/6UCE* targeting near long-range gene targets involves cohesin binding (Cajigas et al., 2018), raising the possibility of roles of additional *Evf2*-RNPs in topological control. X-ChIPseq analysis of *Evf2-*regulated Sox2 and Smc3 binding across *chr6* identifies antagonistic sites of *Evf2* positively (+) regulated Sox2 binding and *Evf2* negatively (-) regulated Smc3 binding (Fig S2A). In order to explore the possibility that Sox2 contributes to *Dlx5/6UCE* targeting, we used chromosome conformation capture (4Cseq) to compare *Dlx5/6UCE* interaction (*Dlx5/6UCEin*) profiles across *chr6* in the presence (Sox2^fl/fl;^*Dlx5/6*cre-) and absence (Sox2^fl/fl;^*Dlx5/6*cre+) of Sox2 in E13.5GE GABAergic progenitors. A subset of *Evf2* regulated *Dlx5/6UCEins* across *chr6* reported in Cajigas et al. (2018) overlap with Sox2 regulated *Dlx5/6UCEins* (Fig S2B-D). *Evf2* and Sox2 shift *Dlx5/6UCE* towards the *5’* end of the repressed target gene *Rbm28* (Fig 2A, red and green arrows, star), overlapping with *Evf2*-5’ recruitment of Sox2 and Smarcc2 (Fig 2B). Sox2 and Smarcc2 binding at the *Rbm28*-5’-*Dlx5/6UCEin* is rescued in *Evf1^TS/TS^* (Fig 2B), linking these events to gene repression. In contrast, Sox2 does not regulate *Dlx5/6UCE* interaction 3’ of the long-range repressed target gene Akr1b8 (Fig 2B, green arrow at *Akr1b10*). Sox2 binding to the *Akr1b10*-*Dlx5/6UCEin* is detected only when *Evf2* is truncated (in *Evf1^TS/TS^*), indicating that *Evf2-3’* prevents Sox2 binding to this site. Together with *Evf2* recruitment of Smarca4 to the Akr1b10-*Dlx5/6UCEin* that is not rescued in *Evf1^TS/TS^*, topological RNP recruitment near Akr1b8 is decoupled from gene repression. These data support that distinct *Evf2-*Sox2 mechanisms contribute to long-range repression of target genes *Rbm28* and *Akr1b8*.

**Figure 2.**
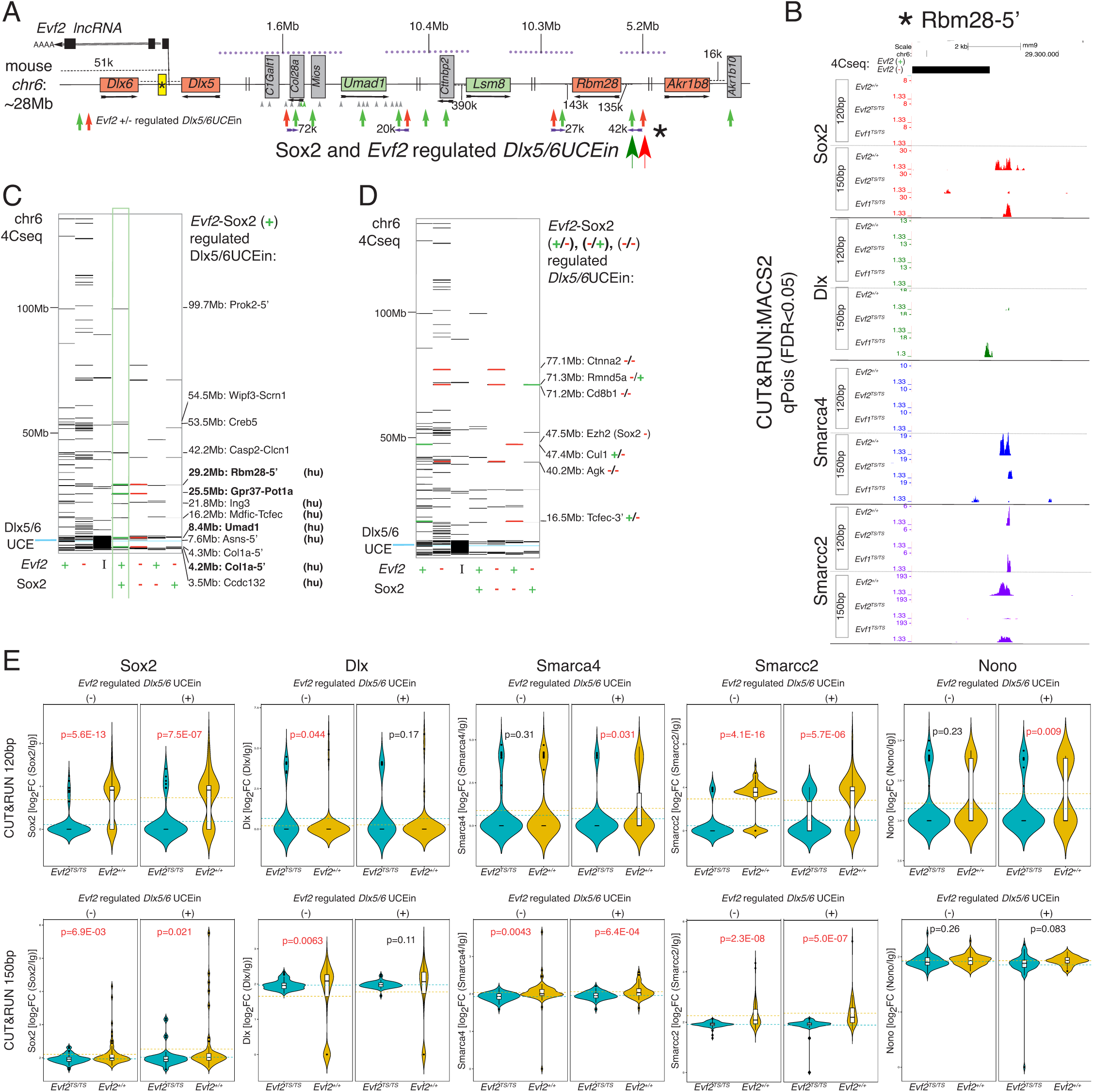
*Evf2* and Sox2 regulated *Dlx5/6*UCEins overlap with *Evf2*-RNP binding on mouse *chr6*. 4Cseq using *Dlx5/6UCE* as bait in E13.5GE from *Sox2^fl/fl^;Dlx5/6cre*+, *Sox2^fl/fl^;Dlx5/6cre*-, *Evf2^+/+^ Evf2^TS/TS^*) and CUT&RUN ChIPseq (E13.5GE from *Evf2^+/+^*, *Evf2^TS/TS,^ Evf1^TS/TS^*) show profiles of *Evf2* regulated *Evf2* RNP binding (Sox2, Dlx, Smarca4, Smarcc2, Nono) across *chr6*. **A.** 4Cseq shows that Sox2 shifts *Dlx5/6UCE* interaction sites (*Dlx5/6UCEins*) towards *Rbm28*-5’ (star, large green arrow), overlapping with *Evf2*-regulated *Dlx5/6UCEin* shift (small green arrow). **B.** CUT&RUN identifies *Evf2-5’* regulation of Sox2 and Smarcc2 binding (absent in *Evf2^TS/TS^*, rescued in *Evf1^TS/TS^*) at the *Rbm28*-5’ *Dlx5/6UCE* targeted site. **C-D.** *Evf2*-Sox2 regulated *Dlx5/6UCEins* across *chr6*. I indicates *Dlx5/6UCEins* that are independent of *Evf2*. Conserved organization of genes on mouse *chr6* and human *chr7* are indicated (hu). 4Cseq analysis using Dlx5/6UCE bait in *Sox^fl/fl^;Dlx5/6cre+/-*, n=4 each genotype. Intersection of two computational methods (FourCseq and DESeq) was used to identify Dlx5/6UCEins (Cajigas et al., 2018). **C**. Positively co-regulated *Dlx5/6UCEin* sites are boxed in green and nearby genes listed. **D.** Genes near antagonistic +/-, -/+ and negatively co-regulated *Dlx5/6UCEin* sites are listed. **E.** Violin Plots comparing *Evf2* regulated RNP binding (CUT&RUN, Sox2, Dlx, Smarca4, Smarcc2, Nono) at *Evf2* positively (+) and negatively (-) regulated *Dlx5/6UCEins* (4Cseq) across *chr6*. Student’s *t*-test, p<0.05 in red, blue (*Evf2^TS/TS^*) yellow (*Evf2^+/+^*). Reads are normalized to IgG.

Across *chr6*, *Evf2* and Sox2 regulated *Dlx5/6UCEins* are categorized into synergistic positive (green +/+, Fig 2C), synergistic negative (red -/-), and antagonistic sites (red/green +/-, -/+) (Fig 2D, Fig S3B-C, Fig S4A). *Evf2*-Sox2 co-regulated *Dlx5/6UCEins* frequently overlap with combinations of *Evf2*-regulated histone marks, and RNP binding (Smc1/3, Sox2, Dlx, Smarca4, Smarcc2, Nono) (Fig S3B-C, Fig S4A). At the *Evf2*-Sox2 positively regulated *Ing3-Dlx5/6UCEin*, *Evf2* regulates H3K4me3, H3K27Ac, and *Evf2*-RNP recruitment (Smc1, Sox2, Dlx, Smarca4 and Smarcc2). Similar to profiles at the *Rbm28-5’ Dlx5/6UCEin* (Fig 2B), Sox2 and Smarcc2 loss in *Evf2^TS/TS^* is rescued in *Evf1^TS/TS^* at the following *Dlx5/6UCEins*: *Ing3*, (Fig S3B), *Ezh2* and *Rmnd5* (Fig S4A), suggesting a common mechanism involving Sox2 and Smarcc2 recruitment.

In order to determine the relationships between *Evf2*-regulated RNP binding (Sox2, Dlx, Smarca4, Smarcc2, Nono), and *Evf2* (+) and (-) regulated *Dlx5/6UCEins* across *chr6,* we combined RNP CUT&RUN results and *Dlx5/6UCE*-4Cseq data from *Evf2^+/+^* and *Evf2^TS/TS^* E13.5GE (Cajigas et al., 2018). Violin plots show that differences between *Evf2*-regulated RNP binding at *Dlx5/6UCEins* are RNP and fragment size - dependent (<120bp, direct binding, >150bp indirect binding) (red p-values indicate significant RNP binding in the presence (yellow) and absence (blue) of Evf2, Fig 2E). *Evf2* increases Sox2 and Smarcc2 binding (120bp and 150bp fragments at both *Evf2* (+) and (-) regulated *Dlx5/6UCEins*; Fig 2E, S4C). Dlx recruitment differs from other RNPs, as *Evf2* decreases binding overall, with significant differences at *Evf2* (-), but not *Evf2* (+) regulated *Dlx5/6UCEins* (Fig 2E). Consistent with site-specific analysis, *Evf1^TS/TS^* rescues a subset of the overall effects, distinguishing roles of the *Evf2-5’* and *-3’* in RNP recruitment at *Dlx5/6UCEins* (Fig S4C).

### *Evf2* regulates Sox2 RNP protein pool targeting to *Dlx5/6* and repressed target genes

We previously found that the *Evf2-5’* (*Evf1^TS/TS^*) is sufficient for both RNA cloud formation and co-localization with *Dlx5/6UCE* (Cajigas et al., 2018). Here we find that *Evf1-3’* RNA continues to be expressed in *Evf2^TS/TS^* nuclei, forming clouds that are properly targeted to repressed genes (*Akr1b8* and *Rbm28*) (Fig 3A-D). These data suggest the *Evf2-5’* and *Evf1-3’ RNA* clouds localize to regulated target genes, but RNA cloud targeting is not sufficient for gene activation or repression.

**Figure 3.**
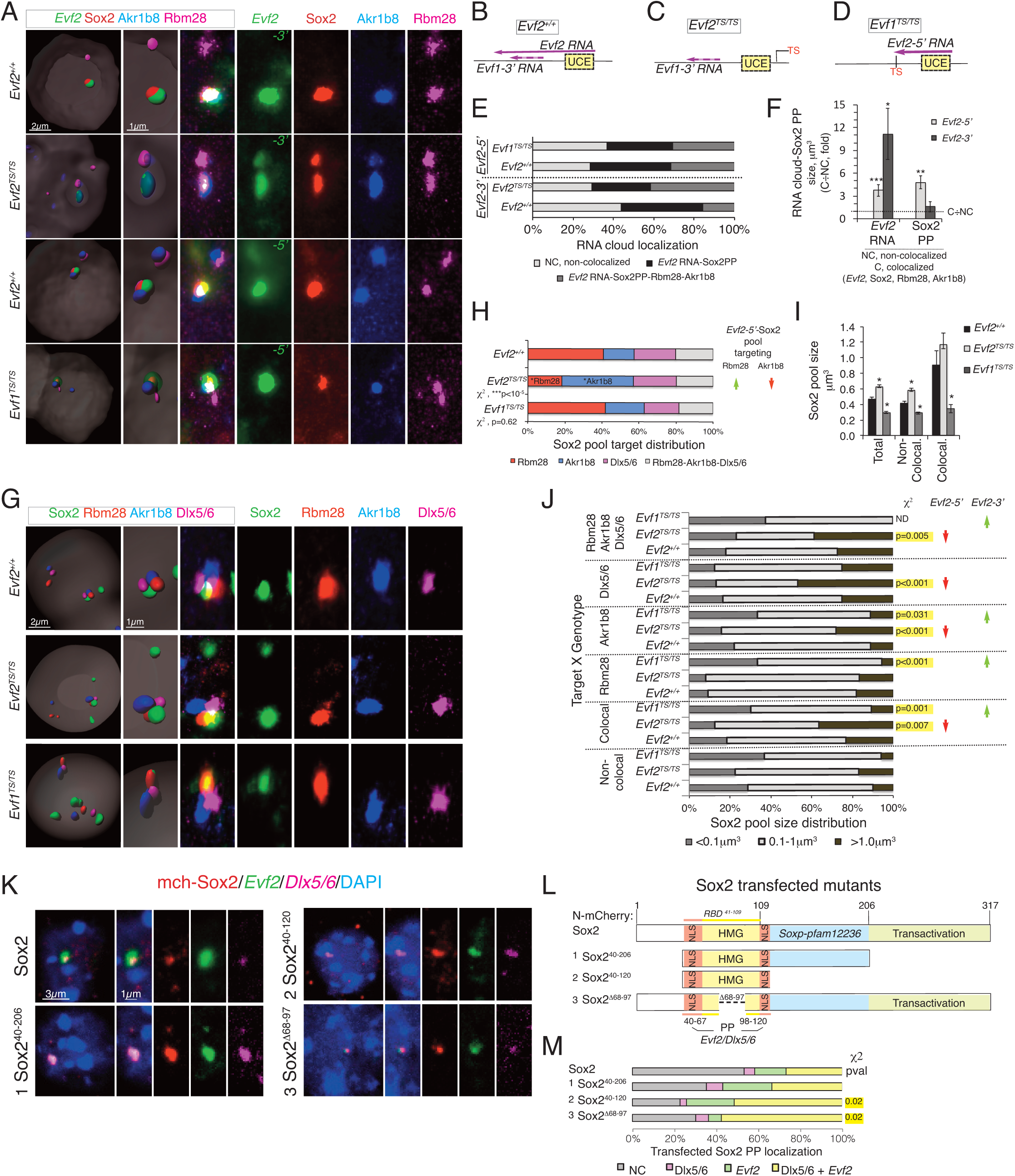
*Evf2* regulates Sox2 protein pool targeting and size. **A**. FISH confocal analysis of *Evf2* RNA clouds (green), Sox2 protein pools PPs (red), *Akr1b8* DNA (blue), *Rbm28* DNA (pink) in *Evf2^+/+^*, *Evf2^TS/TS^*, and *Evf1^TS/TS^* nuclei. Left two panels: IMARIS 3D reconstructions. *Evf2-3’* end or -*5’* end antisense probes differentiate between transcripts. **B-D.** Schematics of *Evf2* full-length, *Evf2* truncated 5’, and *Evf1*-3’ transcripts in different genotypes, TS (triple polyadenylation signal insertion to stop transcription), UCE (*Dlx5/6UCE*). **B**. *Evf2^+/+^* **C**. *Evf2^TS/TS^* **D.** *Evf1^TS/TS^* **E**. Percentages of RNA clouds co-localized with repressed target genes *Akr1b8* and *Rbm28*, or Sox2PP are not significantly changed in *Evf1^TS/TS^* or *Evf2^TS/TS^* mutants. χ2 analysis, p>0.05. n= 144 for *Evf2-5’* probe, n=73 for *Evf2-3’* probe. **F.** In *Evf2^+/+^* nuclei, sizes of *Evf2-5’* and *-3’* RNA clouds co-localized with Sox2, *Rbm28* or *Akr1b8* are significantly larger than non-colocalized clouds; Sox2 protein pool sizes co-localized with *Evf2-5’*, *Rbm28*, or *Akr1b8* are larger than non-colocalized. Student’s t-test (*p<0.05, ***p<0.001), error bars indicate SEM, n=32 (*Evf2^+/+^: Evf2-3’* probe), n=41 (*Evf2^TS/TS^*: *Evf2-3’* probe), n=95. (*Evf2^+/+^: Evf2-5’* probe), n=54 (*Evf1^TS/TS^: Evf2-5’* probe). **G.** Confocal analysis in *Evf2^+/+^*, *Evf2^TS/TS^*, *Evf1^TS/TS^* E13.5 GE nuclei: Sox2 protein pools (green), *Rbm28* DNA (red), *Akr1b8* DNA (blue), *Dlx5/6* DNA (pink). **H.** The number of Sox2 PPs located at *Rbm28*, *Akr1b8*, *Dlx5/6*, and simultaneously at *Rbm28*/*Akr1b8*/*Dlx5/6* in *Evf2^+/+^* (n=65), *Evf2^TS/TS^* (n=88), *Evf1^TS/TS^* (n=56) nuclei were determined; χ2 analysis indicates that Sox2 PP distribution among targets in *Evf2^TS/TS^* (p<10^-5^), but not *Evf1^TS/TS^* differs from *Evf2^+/+^*. *Evf2-5*’, but not 3’, increases the numbers of Sox2 protein pools targeted to *Rbm28* (green arrow), and decreases targeting to *Akr1b8* (red arrow). **I-J.** IMARIS, volumetric measurements of Sox2 PP sizes performed in E13.5 GE nuclei from *Evf2^+/+^*, *Evf2^TS/TS^*, and *Evf1^TS/TS^*. **I**. Sox2 PPs are larger in *Evf2^+/+^* compared to *Evf1^TS/TS^*, but smaller compared to *Evf2^TS/TS^* (total and non-colocalized); colocalization is defined by FISH co-localization with *Rbm28, Akr1b8*, and/or *Dlx5/6UCE*, simultaneously or in any combination), supporting that *Evf2-5’* reduces Sox2 PP size, while *Evf2-3’* increases Sox2 PP size. Sox2 PPs: *Evf2^+/+^* (non-colocalized n= 605, colocalized n=65), *Evf2^TS/TS^* (non-colocalized n= 993, colocalized n=88), *Evf1^TS/TS^* (non-colocalized n= 611, colocalized n=56), ANOVA, post hoc Dunnett’s C test, *p<0.05. **J.** Analysis of Sox2 PP sizes categorized into three sizes: <0.1μm, 0.1μm *-* 1.0μm, >1μm for each genotype and target gene location (non-colocalized, colocalized and specific target gene). Arrows on the right summarize conclusions regarding *Evf2-5’* (red arrows decrease size), and *Evf2-3’* (green arrows increase size) control of Sox2 PP populations. In *Evf2^TS/TS^*, the percentage of Sox2 PPs increases in the >1μm category (colocalized, *Akr1b8* and *Dlx5/6*), whereas profiles are unchanged at *Rbm28*. In *Evf1^TS/TS^* the percentage of Sox2 PPs increase in <0.1μm and/or 0.1μm *-* 1.0μm categories at the expense of larger sizes (>1μm category) at all colocalized targets except *Dlx5/6*. Non-colocalized Sox2 PP size profiles are not altered in *Evf2^TS/TS^* and *Evf1^TS/TS^*. p values from χ2 analysis (compared to *Evf2^+/+^*) are shown on the right. N’s are equivalent as in (I). **K.** FISH confocal analysis of E13.5GE nuclei transfected with mCherry-Sox2 fusion constructs (Sox2 full length, (1) Sox2 40-206, (2) Sox2 40-120, and (3) Sox2 deletion 68-97): mCherry (red), *Evf2* RNA clouds (green), *Dlx5/6UCE* (pink), DAPI (blue). **L**. Schematic of Sox2 mCherry fusion proteins used for transfection into E13.5GE cells. NLS (nuclear localization signal sequences), HMG (high mobility group domain, DNA binding), RBD (RNA binding domain defined from Fig 1), Soxp-pfam12236 (domain shared by Sox protein family members), PP (protein pool). Sox2-HMG domain flanked by NLS is sufficient for Sox2 PP formation, co-localization with *Evf2* RNA clouds, and co-localization with the *Dlx5/6UCE*. RNP co-localization domain is further narrowed to two regions overlapping the NLS (40-67 and 98-120). **M**. Sox2 mutants differ in the percentage of Sox2 PP co-localized with *Evf2* RNA and *Dlx5/6UCE* (yellow bars), supporting that Sox2 67-98 inhibits RNP association. p=0.02 from χ2 analysis are shown on the right (χ2 analysis of distributions), n=60 (mchSox2), n=51 (mchSox2^40-206^ mutant 1), n=35 (mchSox2^40-120^ mutant 2), n=33 (mchSox2^Δ68-97^ mutant 3).

Confocal microscopy in wildtype E13.5GE nuclei previously showed that *Evf2*-RNPs are enriched within *Evf2* RNA clouds (Dlx (Feng et al., 2006), Smarca4 (Cajigas et al., 2015), Smc1 (Cajigas et al., 2018)). Therefore, we used RNA FISH combined with immunofluorescence to study a possible role for *Evf2* in Sox2 localization. Surprisingly, unlike diffusely distributed Smarca4 protein enriched in *Evf2*-RNA clouds (Cajigas et al. 2015), Sox2 forms heterogeneously sized protein pools (PPs) in E13.5GE nuclei (Fig 3A and 3G). Sox2 PPs co-localize with *Evf2* RNA clouds (Fig 3A) and with *Evf2-*repressed target genes, *Akr1b8*, *Rbm28* (Fig 3G). A subset of Sox2 PPs co-localize with *Evf2-5’* and *-3*’ RNA clouds (Fig 3A-F). While the percentages of *Evf2-5’* and *-3’* RNA clouds colocalized with Sox2 PPs is not significantly changed in *Evf1^TS/TS^* and *Evf2^TS/TS^* nuclei, the sizes of *Evf2-5’* and *-3’* RNA clouds are significantly larger when colocalized with Sox2-PPs or *Evf2*-repressed target genes (Fig 3F). In addition, *Evf2-5’* associated Sox2 PPs colocalized with repressed target genes are larger than non-colocalized (Fig 3F). Visualization of Sox2 PPs in *Evf1^TS/TS^* and *Evf2^TS/TS^* nuclei indicates that Sox2 PPs continue to be targeted to *Rbm28*, *Akr1b8* and *Dlx5/6* (Fig 3G). However, in *Evf2^TS/TS^* the percentage of Sox2-PPs colocalized with *Akr1b8* increases at the expense of *Rbm28*, a shift that is rescued in *Evf1^TS/TS^* (Fig 3H). Total and non-colocalized Sox2-PPs are larger in *Evf2^TS/TS^*, and smaller in *Evf1^TS/TS^* nuclei (Fig 3I), where non-colocalization is defined as non-overlapping with *Rbm28*, *Akr1b8*, or *Dlx5/6*. Increased variability of *Evf2^TS/TS^* colocalized Sox2-PPs (Fig 3I) led to binning Sox2-PPs into three size groups (<0.1µm^3^, 0.1-1µm^3^, >1µm^3^) for comparisons of size distributions at specific targets in *Evf2^+/+^*, *Evf2^TS/TS^* and *Evf1^TS/TS^* nuclei (Fig 3J). Co-localized Sox2PP sizes are increased in *Evf2^TS/TS^* (red arrows) and decreased in *Ev1^TS/TS^* (green arrows), with larger effects observed at *Akr1b8* (alone or complexed with *Rbm28* and *Dlx5/6*) (Fig 3J). At *Dlx5/6*, the percentage of *Evf2^TS/TS^* Sox2-PPs >1µm^3^ increases at the expense of 0.1-1µm^3^ (Fig 3J, red arrow). This shift in Sox2PP sizes is rescued in *Evf1^TS/TS^*, supporting the significance of Sox2PP size regulation at *Dlx5/6UCE* in gene repression. Together, these data support that *Evf2-5’* and *-3’* differentially regulate Sox2PP gene targeting and size distributions in a site-specific manner.

We next defined Sox2 protein domains involved in Sox2PP association with *Evf2* RNA clouds and *Dlx5/6*, using FISH analysis of N-terminally tagged, mcherry-Sox2 (mch-Sox2) transfected into E13.5GE’s. Transfected Sox2 forms PPs that colocalize with endogenous *Evf2* RNA clouds and/or *Dlx5/6* (Fig 3K-L). Sox2^40-120^ (mutant 2), a domain involved in RNA binding (Fig 1G) is sufficient for PP formation, and for co-localization with the *Evf2* RNA cloud and *Dlx5/6* (Fig 3K). However, deletion of Sox2^66-97^ increases co-localization with *Evf2* and *Dlx5/6*, supporting that Sox2 HMG internal amino acids inhibit *Evf2*-RNP association. This mutational analysis indicates sufficiency of amino acids within the Sox2 HMG domain (Sox2^40-67^ and Sox2^98-120^) for co-localization with *Evf2 and Dlx5/6*. It is possible that amino acids within the NLSs contribute to Sox2 PP formation or co-localization with *Evf2* and *Dlx5/6*. However, the NLS sequences were not mutated in this analysis, as these mutations would interfere with nuclear localization.

### *Evf2* regulates targeting and sizes of Sox2 associated PP forming *Evf2*-RNPs

*Evf2* RNA cloud co-localization with Sox2 PPs raised the possibility that other *Evf2*-RNPs also form PPs, leading to a screen for *Evf2* RNPs that colocalize with Sox2 PPs. In E13.5GE nuclei, we identify *Evf2* RNPs Nono, Smc1a, Smc3 and Smarcc2 PPs that colocalize with Sox2 PPs (Fig 4A), *Evf2* repressed genes (*Rbm28* and *Akr1b8*), and *Dlx5/6* (Fig 4B). The percentage of Smarcc2 and Nono PPs co-localized with *Rbm28*/*Akr1b8*/*Dlx5/6* decreases in *Evf2^TS/TS^*, while Smc3 PP co-localization increases in *Evf1^TS/TS^*, indicating distinct roles of *Evf2-5’* and *-3’* ends in *Evf2*-RNP targeting (Fig 4C). Analysis of PPs shows RNA-dependent regulation of RNP PPs size. Analysis of averaged non-colocalized Smarcc2, Smc1a and Smc3 PP sizes shows increases in *Evf2^TS/TS^* nuclei compared to *Evf2^+/+^*, while Smarcc2 and Smc1a PPs sizes are rescued in *Evf1^TS/TS^*, and Smc3 PPs sizes are not (Fig 4D). Similar to Sox2, colocalized Smarcc2 and Smc1a PPs in *Evf2^TS/TS^* show high degrees of variation in sizes (Fig 4D). Further analysis by binning *Evf2*-RNP-PPs into three sizes indicates that, in *Evf2^TS/TS^*, the percentage of Smarcc2 PPs in the largest size group (>0.3µm^3^) is significantly increased, at the expense of the smallest size group (<0.1µm^3^); χ2 analysis on *Evf2^TS/TS^* colocalized Smarcc2 PPs is not performed due to small numbers (Fig 4E). In the case of Nono-PPs, binning reveals an increase in 0.2-0.5µm^2^ in *Evf2^TS/TS^* rescued in *Evf1^TS/TS^* (Fig 4E). Thus, *Evf2*-5’ decreases the sizes of all colocalized *Evf2*-RNP PPs tested (Fig 4E), including Sox2 (Fig 3J), while *Evf2-5’* and *- 3*’ differential control of *Evf2*-RNP targeting is site specific (Sox2), and RNP-specific.

**Figure 4.**
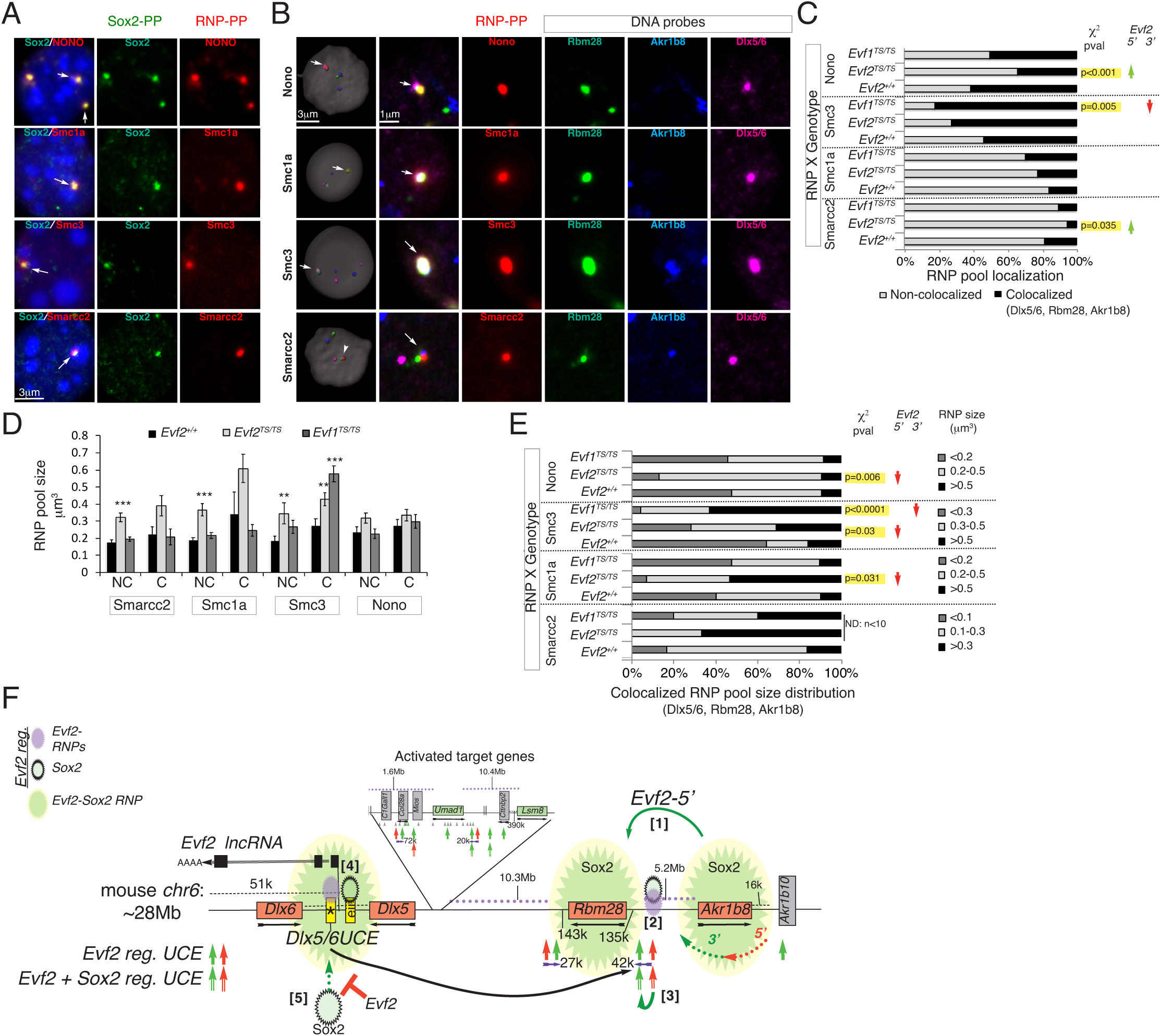
*Evf2* regulates targeting and sizes of Sox2 co-localized PP forming *Evf2*-RNPs (Nono, Smc1a, Smc3 and Smarcc2). **A.** *Evf2*-RNPs (Nono, Smc1a, Smc3 and Smarcc2) form PPs that co-localize with Sox2 PPs in *Evf2^+/+^* E13.5GE nuclei (confocal analysis of Sox2 protein (green), and specific RNPs (Nono, Smc1a, Smc3, Smarcc2, red). **B**. FISH confocal analysis of *Evf2*-RNPs (Nono, Smc1a, Smc3 and Smarcc2) forming PPs (red) that co-localize with *Evf2* repressed target genes *Rbm28* (green) and *Akr1b8* (blue), and *Dlx5/6UCE* (pink). **C**. *Evf2-5’* and *-3’* regulation of *Evf2*-RNP (Nono, Smc1a, Smc3 and Smarcc2) co-localization with *Rbm28*, *Akr1b8* and *Dlx5/6UCE*. p values from χ2 analysis are shown on the right. Green arrows indicate that *Evf2-5’* increases co-localization of Smarcc2 and Nono, whereas *Evf2-3’* decreases co-localization of Smc3. C-E. n’s for PPs: *Evf2^+/+^* (Smarcc2 n= 64, Smc1a n= 55, Smc3 n=46, Nono n=68), *Evf2*^TS/TS^ (Smarcc2 n= 53, Smc1a n= 66, Smc3 n=44, Nono n=90), *Evf1^TS/TS^* (Smarcc2 n= 55, Smc1a n= 60, Smc3 n=40, Nono n=69). **D**. *Evf2-5’* and *-3’* regulation of non-co-localized and co-localized (*Akr1b8*, *Rbm28*, *Dlx5/6UCE*) *Evf2*-RNP (Nono, Smc1a, Smc3 and Smarcc2) PP sizes, Student’s t-test, pairwise comparisons with *Evf2^+/+^*, *p<0.05, ***, p<0.001. **E.** *Evf2* regulation of *Evf2*-RNP-PP (Smc1a, Smc3, Nono) sizes at co-localized sites (Dlx5/6, Rbm28, Akr1b8). The profiles of *Evf2*-RNP PP sizes are compared in *Evf2^+/+^*, *Evf2^TS/TS^* and *Evf1^TS/TS^* E13.5 GE; p values based on χ2 analysis of size distributions are shown on the right. PP sizes are binned in optimal ranges for each RNP (shown on the right). *Evf2-5’* decreases sizes of Smc1a, Smc3 and Nono PPs (red arrows). In *Evf2^TS/TS^*, Smarcc2 PPs <0.1 μm^3^ are not detected; in addition, n <10 for both *Evf2^TS/TS^* and *Evf1^TS/TS^*, and therefore, DD was not calculated for Smarcc2 PPs (ND). *Evf2-3’* decreases Smc3 co-localized PP sizes (red arrow). **F.** Modelling multi-step contributions of *Evf2*-Sox2 interactions in *Dlx5/6UCE* targeting and activity during gene repression, described in detail in the Discussion.

## Discussion

Understanding lncRNA-dependent chromosome topological control requires mechanistic experiments that define individual contributions of lncRNA-RNPs. Many methods have been developed to identify regulatory lncRNA-RNP complexes, with the most recent utilizing CRISPR technology (Yi et al., 2020). The *Xist-RNP* is among the most well-characterized lncRNA regulatory complexes, with comprehensive lists of *Xist*-RNPs from mass spectrometry sequencing, as well as validated functions of individual proteins (Chen et al., 2016; Chu et al., 2015; Dossin et al., 2020; Minajigi et al., 2015; Yi et al., 2020). In addition to RNA cloud formation, *Evf2* and *Xist* share functional characteristics (gene repression, chromatin remodeling effects, topological control) (Giorgetti et al., 2016; Jegu et al., 2019; Nora et al., 2012), and formation of a similar sized RNP (*Xist*-RNP^85^ (Chu et al., 2015), where 16/85 proteins are shared with *Evf2-RNP*). In a separate characterization of the *Xist*-RNP, Smarca4, the catalytic subunit of SWI/SNF-related chromatin-remodeling complex, and cohesin (Smc1 and Smc3) were identified, increasing similarities between *Evf2* and *Xist* (Minajigi et al., 2015). *Evf2* RNA-Smarca4 interactions inhibit chromatin remodeling activity through inhibition of ATPase activity (Cajigas et al., 2015), a mechanism also shared during *X-*inactivation (Jegu et al., 2019). Furthermore, *Evf2*-dependent recruitment of Smc1 and Smc3 at key topologically controlled sites, supports that *Evf2* targets *Dlx5/6UCE* to long-range target genes by regulating Smc1 and Smc3 binding (Cajigas et al., 2018). LncRNA-dependent cohesin recruitment is also shared with *Xist* (Minajigi et al., 2015) and ThymoD lncRNA (Isoda et al., 2017), raising the possibility that topological control is a shared function of cloud forming lncRNAs. Given that components of the *Evf2*-RNP interact with other lncRNAs, the study of gene regulation by the *Evf2*-RNP provides important insight into general mechanisms of gene regulation by lncRNAs.

While Sox2 transcriptional activities have been extensively characterized, new roles are emerging for Sox2 RNA binding activities. The *Evf2* RNP Sox2 HMG contains both DNA and RNA binding domains (Holmes et al., 2020), necessary for *Evf2* binding (Fig 1E) and sufficient for PP localization in vivo (Fig 3K,L). Together with genetic epistasis experiments (Fig 1H), these experiments support that *Evf2* gene repression occurs through direct antagonism in which *Evf2* lncRNA binds to Sox2, reduces the binding of Sox2 to DNA regulatory elements, thereby decreasing enhancer activity.

However, our evidence also supports a role for Sox2-lncRNA interactions in regulating enhancer targeting, and reciprocal effects of *Evf2* on Sox2 protein recruitment, in events that likely precede direct effects on enhancer activity (Fig 4F). Key to this multi-step model of regulation is formation of the *Evf2-*RNP, assembled on both lncRNA and protein scaffolds. LncRNA scaffolds bridge individual RNA binding proteins through promiscuous RNA binding properties, as shown in this report for Sox2 (*Evf2*-Sox2 (Fig 1E)) and previous work for chromatin remodelers (*Evf2*-Smarca4, *Evf2*-Smarcc2, *Evf2*-Smarcc1 (Cajigas et al., 2015)) (Fig 1A). Multivalent proteins bridge non-RNA binding RNPs (Dlx, Smarcb1) to the lncRNA through interactions with promiscuously bound RNA binding proteins Sox2 (Fig 1B) and Smarca4 (Cajigas et al., 2015). High-affinity promiscuous RNA binding properties, reported for PRC2 (Davidovich et al., 2013), Smarca4 (Cajigas et al., 2015), and Sox2 (this report and (Holmes et al., 2020)), explain the ability of the lncRNA to act as an organizer, with “glue” like properties, controlling the availability of specific RNPs in a site-specific manner. In such a model, DNA site specificity is determined by enhancer transcription and sequence: **a**) enhancer transcription produces a lncRNA that is retained and provides a scaffold where the RNP grows, and **b),** enhancer sequences recruit and stabilize TF binding (Dlx and Sox2). *Evf2* RNA clouds and *Evf2-*RNP-Sox2 PP sizes are larger when colocalized with *Dlx5/6UCE* and/or specific DNA target genes than non-localized (Fig 2F, I), supporting the idea that the *Evf2*-RNP grows at key DNA regulatory sites.

The model in Fig 4F schematizes *Evf2-Sox2* interactions involved in multiple steps of gene repression. In [1], *Evf2-5’* limits the number of *Evf2*-Sox2 RNPs targeted to *Akr1b8*, by shifting *Evf2*-Sox2 RNPs towards *Rbm28*, as supported by confocal FISH analysis (Fig 2H). In addition, increased Sox2-PPs targeting to *Akr1b8* increases Sox2-PP sizes in *Evf2^TS/TS^*, supporting the idea that *Evf2-5’* repression of *Akr1b8* occurs by limiting the availability of Sox2 (an activator of *Dlx5/6UCE*) through targeting and size regulation, thereby sequestering Sox2 PPs within the *Evf2*-RNP. Furthermore, *Evf2-5’* (sufficient for *Akr1b8* repression) properly balances the numbers of *Evf2*-Sox2RNPs at *Akr1b8* and *Rbm28* in *Evf1^TS/TS^*, linking rescue of *Evf2*-Sox2RNP targeting to rescue of gene repression.

In Fig 4F[2], *Evf2-5’* recruits *Evf2*-RNPs Sox2 and Smarcc2 to the *Rbm28-5’-Dlx5/6UCEin,* as supported by CUT&RUN binding profiles from *Evf2^+/+^*, *Evf2^TS/TS^* and *Evf1^TS/TS^* (Fig 2B). *Evf2* regulation of histone methylation (H3K4me3/H3K37me3), histone acetylation (H2K27ac) and Smc1 binding at *Rbm28-5’-Dlx5/6UCEin* supports concerted actions (Fig S3B, (Cajigas et al., 2018)), increasing the significance of this site. Overlapping *Evf2-*Sox2 regulated *Dlx5/6UCEins* map across *chr6*, a subset located within 50kb of *Evf2* RNP recruited sites and histone modifications, notably: the 16.2Mb Mdfic-Tcfec gene desert [*Evf2-5’* regulated Sox2, Smarcc2], Ing3 [Sox, and *Evf2-5’* regulated Dlx, Smarca4, Smarcc2], Prok2 [*Evf2-5’* Smarcc2 regulated], Rmnd5a [Dlx, and *Evf2-5’* regulated Sox2, Smarca4, Smarcc2], (Fig S3B, C, S4A), revealing the combinatoric nature of *Evf2* regulation. Interestingly, Sox2 negative regulation of *Dlx5/6UCEin* at the *Ezh2-5’* promoter overlaps with *Evf2* (+) regulated Sox2, Smarca4, Smarcc2, Nono, and (-) regulated Dlx, suggesting topological control of a key epigenetic regulator (Fig S4A). Recently, Ezh2 inhibitors have been approved to treat cancer (Richart and Margueron, 2020), but also potentially increase seizure susceptibility in mice (Wang et al., 2020). While our studies have focused on the *Evf2*-regulated GRN in embryonic GEs, it will be important to determine whether *Evf2*-regulation of RNPs, in combination with Sox2 controlled *Dlx5/6UCEin* at the *Ezh2-5’* contribute to seizure susceptibility phenotype observed in mice lacking *Evf2* (Cajigas et al., 2018). Consistent with our previous reports of *Evf2* topological and histone modification control (Cajigas et al., 2018), *Evf2* regulated RNP binding sites and *Evf2*-Sox2 regulated *Dlx5/6UCEins* are also detected outside the *Evf2* 27Mb GRN region, revealing chromosome-wide effects.

In Fig 4F[3], *Evf2* and Sox2 shift *Dlx5/6UCE* towards the *Rbm28-5’* promoter, as supported by combined 4Cseq analysis of *Evf2^+/+^*, *Evf2^TS/TS^* (Cajigas et al., 2018), and *Sox2^fl/fl^;Dlx5/6cre+/-* genetic models (Fig 2A, S2B-D, S3B). One caveat to *Evf2* gene repression through limiting Sox2 PP activator is that in *Evf2^TS/TS^*, the numbers of Sox2- PPs targeted to *Rbm28* (also a repressed target gene) decreases, while Rbm28 gene expression increases. One possibility is that the extent of repression is affected by a combination of Sox2 targeting and size regulation at the two repressed target genes: loss of *Evf2* causes ∼15-fold increase in *Akr1b8*, but ∼2-fold increase in *Rbm28*. Another possibility is that mechanisms of *Evf2*-Sox2 antagonism differ between *Akr1b8* and *Rbm28*, as reflected by target gene-dependence. This is supported by 4Cseq analysis showing that Sox2 shifts *Dlx5/6UCE* towards *Rbm28-5’* (overlapping with *Evf2*-*Dlx5/6UCE* regulation), but does not affect *Dlx5/6UCEin* near *Akr1b8*. Site-specific *Evf2*-Sox2 synergistic and antagonistic regulation of *Dlx5/6UCEins* across *chr6* further supports variable combinations of events during topological control.

In Fig 4F[4], *Evf2* recruits RNPs to *Dlx5/6* intergenic enhancers *Dlx5/6UCE* and *Dlx5/6eii*, consistent with previous reports of *Evf2* regulated Dlx and Smarca4 recruitment (Bond et al., 2009; Cajigas et al., 2015). Although *Dlx5/6eii* lacks ultraconserved sequences, *Dlx5/6* intergenic enhancers are functionally similar, both regulated by Dlx and *Evf2* (Feng et al., 2006; Zerucha et al., 2000). Deletion of *Dlx5/6eii* in mice alters gene expression in developing and adult interneurons (Fazel Darbandi et al., 2016), supporting both overlapping and distinct functions in vivo. A key distinguishing feature of *Dlx5/6UCE* is transcription into a spliced, polyadenylated lncRNA (*Evf2*), whereas stable transcripts from *Dlx5/6eii* are not detected. With respect to *Evf2-*RNP binding, Sox2 binds both enhancers, while Dlx, Smarca4 and Smarcc2 binding is limited to *Dlx5/6UCE*. *Evf2-*RNP recruitment also differs: *Evf2* regulates Sox2 binding to *Dlx5/6eii*, but not to *Dlx5/6UCE*, whereas *Evf2* regulates Dlx, Smarca4, and Smarcc2 binding to *Dlx5/6UCE,* but not to *Dlx5/6eii*. Such differential recruitment is consistent with a “separation of inputs” hypothesis that enables shadow enhancer pairs to buffer noise better than duplicate enhancers (Waymack et al., 2020). Thus, *Evf2* recruitment of Sox2 to *Dlx5/6eii* may reflect the need to precisely regulate levels of Sox2, beyond that necessary for factors that are recruited to *Dlx5/6UCE*, where on/off decisions are made.

In Fig 4F[5], Sox2 increases *Dlx5/6UCE* activity, while *Evf2* decreases *Dlx5/6UCE* activity by directly binding to Sox2-HMG DNA binding domain, antagonizing activation. This model is supported by REMSAs (Fig 1E-G), and luciferase reporter assays (Fig 1I). Evidence showing that the Sox2 HMG-containing (and adjacent NLS) domain is sufficient for Sox2PP formation and localization to *Evf2* RNA clouds and *Dlx5/6UCE*, supports the significance of *Evf2*-Sox2 HMG interactions, in vivo. The ability of *Evf2* lncRNA to directly inhibit chromatin remodeling by inhibiting ATPase activity (Cajigas et al., 2015), and extension of this mechanism to X-inactivation (Jegu et al., 2019), adds a 6^th^ dimension to lncRNA-mediated transcriptional regulation that is not schematized in the model.

Together, these data suggest that *Evf2* gene repression involves multiple levels of *Evf2*-RNP regulation, including PP targeting and size regulation, site-specific recruitment and sequestration that ultimately affect both *Dlx5/6UCE* targeting and activity.

How individual *Evf2*-RNPs regulate *Evf2* RNA cloud assembly at specific DNA sites is not known. Given that Nono has been shown to assemble the NEAT lncRNA into paraspeckles, it is possible that Nono plays a similar role in the *Evf2*-RNP (Yamazaki et al., 2018). Experiments that tag lncRNA clouds and follow assembly through high-resolution microscopy will be important in validating static studies. Since only 1-2 *Evf2* RNA clouds are detected in the nucleus, either the *Evf2*-RNP moves with *Dlx5/6UCE* to repressed target genes, or DNA looping (Rao et al., 2017) brings *Evf2*-RNP bound *Dlx5/6UCE* closer to target genes (Cajigas et al., 2018). Analysis of additional PP forming *Evf2-*RNPs (Smarcc2, Smc1a/3 and Nono) indicates that Sox2 PP size regulation and targeting is shared with other *Evf2-*RNP-PPs. As detected for Sox2 PPs, *Evf2-5’* decreases PP sizes formed by Nono, Smc3, Smc1a, and Smarcc2. However, *Evf2-3’* increases Sox2-PP sizes, while decreasing Smc3-PP sizes, supporting differential roles of the *Evf2* RNA in RNP-PP growth, and raising the possibility of *Evf2*-RNP polarity.

Experiments in this report reveal a complex, multi-step role for *Evf2*-Sox2 interactions in enhancer targeting and direct antagonism of *Dlx5/6UCE* activity during gene repression. RNA cloud regulation of PP forming RNPs. How each of the individual *Evf2* RNPs (of the 87 identified) contributes to *Dlx5/6UCE* targeting and activity regulation, the relationship of these functions to RNP assembly, and to common mechanisms of lncRNA transcriptional control remain open questions. Our data suggests that by coupling recruitment and sequestration of critical RNP proteins at key steps of enhancer targeting and activity, *Evf2* RNPs selectively regulate genes across megabase distances. We propose that *Evf2* RNA clouds function as both chromosome and protein organizers, contributing to dynamic regulation at enhancers.

## Materials and Methods

### Recombinant protein pulldown

Co-immunoprecipitation experiments on 6X-his tagged proteins purified from E.coli or flag-tagged baculovirus proteins was performed as described (Cajigas et al., 2015).

### RNA Electrophoretic Mobility Assay (REMSA)

The NIR probe was generated by in vitro transcription of pGEM-Evf2(UCR), and REMSAs performed as described (Cajigas et al., 2015).

### Primary embryonic brain MGE transfections

For all transfections, MGE tissues were dissected from E13.5 Swiss Webster embryos. Luciferase assays were performed as described (Cajigas et al., 2018).

### Combined DNA-RNA FISH with Immunofluorescence

DNA-RNA FISH was performed as described (Cajigas et al., 2018). The templates for the nick translation reactions were obtained from the BACPAC Resources Center (Children’s Hospital Oakland Research Institute): Dlx5/6 region: WI1-1693G2, Akr1b8 region: RP23-120B14, Rbm28 region: RP23-276H18. The digoxigenin labeled RNA probe was generated as described previously (Feng et al., 2006).

### Cut& Run

Cut&Run was performed as previously described (Skene et al., 2018) with some modifications. The number of samples was determined according to the number of antibodies to be tested and the number of replicates (2 replicates per antibody and 2 replicates for IgG). The appropriate volume of Concanavalin A magnetic beads (10 μl per sample) were mixed into 1.5 ml of ice-cold Binding Buffer (20 mM HEPES-KOH pH 7.9, 10 mM KCl, 1mM CaCl_2_, 1mM MnCl_2_) and placed on a magnetic stand for 2 min. The supernatant was removed, and the beads were washed with 1.5 ml Binding Buffer. After magnetic separation of the beads, the supernatant was removed, and a volume of Binding Buffer equal to the initial beads volume (10 μl per sample) was added to the beads. The beads were placed on ice. E13.5 ganglionic eminences were isolated from embryos in L15 medium (6 Embryos per genotype). Tissues were pooled for each genotype, triturated by pipetting, and filtered through a cell-strainer capped 5 ml polystyrene round-bottom tube (BD Falcon) to generate single-cell suspensions. pA-Mnase was optimized and used at 700 ng/ml in Digitonin Buffer and 2 pg/ml heterologous spike-in DNA was used. Cut&Run libraries were prepared using the KAPA Hyper Prep Kit protocol, and quality was analyzed by TapeStation prior to sequencing on a NovaSeq 6000 (SP 100 cycles).

### Cut&Run data processing

Paired-end Cut&Run reads of different transcription factors and their respective Ig-controls from Evf2+/+ and Evf2TS/TS Dlx5/6UCE samples were first mapped on mm9 genome using Bowtie2 v2.1.0 (Langmead and Salzberg, 2012) with options ‘--local --very-sensitive-local --no-unal --no-mixed --no-discordant --phred33 -I 10 -X 700’. We used the Picard toolkit command ‘MarkDuplicates’ to mark PCR duplicates and remove them from the final mm9 genome mapped bam files. Next, we separated the sequence fragments into ≤ 120 and ≥ 150-bp size classes that provided the mapping of the local vicinity of a DNA-binding protein. The base-pair sizes can vary depending on the steric access to the DNA by the tethered MNase (Skene et al., 2018). Fragments mapping to repeat elements were removed, and replicates were joined before peak calling. The peak calling was performed using MACS2 (Zhang et al., 2008) callpeak options ‘-t -c –f BED -g mm --keep-dup all --bdg –nomodel --slocal 500 --llocal 5000 –-extsize 120/150’. An FDR cutoff of 0.05 was used to call the final set of peaks (Janssens et al., 2018). Differential Cut&Run analysis of transcription factor binding peaks between two Dlx5/6UCE conditions was performed using MACS2 program by treating one of the samples as the “control” for the other.

### 4C-Seq

4C was performed as previously described (van de Werken et al., 2012), with some modifications. GEs from Sox2 ^fl/fl^ Cre+ and Sox2 ^fl/fl^ Cre-single embryos were dissected in L15 and kept in separate tubes, and processed as described (Cajigas et al., 2018).

### Dlx5/6UCE 4C-seq differential data analysis

4C methods were performed as described (Cajigas et al., 2018)using Bowtie2 v2.1.0 (Langmead and Salzberg, 2012), DESeq2 (Love et al., 2014), FourCSeq program (Klein et al., 2015). To avoid method-specific biases, interaction sites that were assigned the same label (+, -, I) by the two different 24 approaches (DESeq2 and FourCSeq) were called 4Cseq-intersectional computational method sites (4Cseq-ICMs). For Sox2flfl +/cre 4C-seq differential data analysis, We applied FourCSeq on our *Sox2flfl +* and *Sox2flfl cre* 4Cseq samples and retrieved Sox2 regulated 4C interactions (p-adjusted value < 0.05 and an absolute log2 fold change ≥ 2). To avoid method-specific biases, we retained a common set of 244 4C differential sites.

### Dlx5/6UCE and Sox2flfl +/cre site overlap

Evf2+/+ and Evf2TS/TS Dlx5/6UCE and *Sox2flfl* +/cre 4C-peak overlap was performed using ‘bedtools window’ function (Quinlan and Hall 2010) with a window span of 50kb.

### Dlx5/6UCE 4C-seq and Cut&Run signal overlap

*Evf2^+/+^* and *Evf2^TS/TS^ Dlx5/6UCE* 4C-seq peaks were first mapped on Cut&Run differential transcription factor peaks using bedtools ‘intersect’ function (Quinlan and Hall, 2010). The overlapping set of 4C-seq peaks from a *Dlx5/6UCE* condition were then mapped on the respective Cut&Run Ig-normalized transcription factor signal data. The log2 fold-enrichment of Ig-normalized signal was generated using MACS2 ‘bdgcmp’ command (Zhang et al., 2008). The violin plots were made using ‘ggpubr’ R package (Kassambara).

## Author contributions

Conceptualization, I.C., J.D.K. Methodology, I.C., J.D.K, Software, A.C., F.A., Validation, I.C., M.L., M.B., L.C., Formal Analysis, J.D.K., F.A., A.C., Investigation, I.C.,M.L., K.R.S., M.B., L.C., H.L., Data Curation, J.D.K., A.C., Writing-original draft, J.D.K., Writing review&editing, J.D.K., I.C., F.A., Visualization, J.D.K., Supervision, J.D.K., Project Admin. J.D.K., Funding acquisition, J.D.K., and F.A

## Acknowledgments

We thank R. Kingston and J. Cochrane (Harvard) for flag-tagged Smarca4 protein, funding NIGMS R35GM128938 (F.A.), NIMH R01MH111267 (J.D.K). Authors have no conflicts of interest.

## Supplemental Information

### RESOURCE AVAILABILITY

#### Lead Contact

Further information and requests for resources and reagents should be directed to Jhumku D. Kohtz.

#### Materials Availability

Plasmids and mice generated in this study will be available upon request.

#### Data and Code Availability

- The Sox2flfl +/cre 4C-seq data generated and analyzed during this study are available at our local repository https://informaticsdata.liai.org/Collaborators/northwestern/Sox2_Paper_Data/4C_Data [prior to data deposition in NCBI, password will be available upon request]; <u>the data will be deposited in a public repository (NCBI).</u>
- The Cut&Run data generated and analyzed during this study are available at our local repository https://informaticsdata.liai.org/Collaborators/northwestern/Sox2_Paper_Data/CutRun_Data [prior to data deposition in NCBI, password available upon request]; <u>the data will be deposited in a public repository (NCBI).</u>
- The published article Cajigas et la., Molecular Cell, 2018 (https://doi.org/10.1016/j.molcel.2018.07.024) includes the DLX5/6UCE Chip-Seq, 4C-seq dataset (GSE117184) generated and analyzed during this study.
- Original confocal images have been uploaded to Mendeley (Confocal Images : Kohtz, Jhumku (2020), “Image data for Cajigas et al 2020”, Mendeley Data, V1, doi:

### EXPERIMENTAL MODEL AND SUBJECT DETAILS

*Animal models: Evf2^TS/TS^* (Bond et al. 2009) and *Evf1^TS/TS^* (Cajigas et al. 2018) were crossed to C57/Bl6 for one generation, and maintained on a mixed background (C57/Bl6, 129Sv, FVB). Sox2^fl^ Dlx5/6 Cre: Mice containing Lox P sites flanking Sox2 (Sox2^fl.^ Jackson Labs) were crossed to a mouse strain that expresses the Cre recombinase under the control of Dlx5/6 regulatory sequences (Dlx5/6cre, Jackson Labs), for the conditional removal of Sox2 in an interneuron cell population. These mice were maintained on a mixed background (C57/Bl6, 129Sv, CD1).

### METHOD DETAILS

#### Recombinant protein pulldown

Recombinant proteins were purified from bacteria utilizing standard methods. Proteins were incubated in 200 μl NETN Buffer (100 mM NaCl, 0.5 mM EDTA, 20 mM Tris-HCl, pH 8.0, 0.5 % NP-40) for 1 hour at 4°C with rotation. 30 μl of Glutathione agarose beads were washed with NETN buffer and added to each sample. Samples were incubated for 1 hour at 4°C with rotation. Beads were pelleted by centrifugation and washed 3 times with 1 ml NETN buffer and one time with 1x PBS. Proteins were eluted by adding Protein loading buffer (62.5 mM Tris, pH 6.8 10% Glycerol, 2% SDS, 5% B-mercaptoethanol, 0.002% Bromophenol blue) and incubating at 95°C for 5 min. Samples were analyzed by Western Blot.

#### RNA Electrophoretic Mobility Assay (REMSA)

The NIR probe was generated by in vitro transcription of *pGEM-Evf2(UCR)* (Cajigas et al. 2015), a plasmid containing 115 nt of *Evf2*, including the ultraconserved sequence. 1 μg of SalI linearized DNA template, 5 mM DTT, 0.6 μl RNasin (Promega), 0.5 mM ATP, 0.5 mM CTP, 0.5 mM GTP, 12.5 μM UTP, 20 mM Aminoallyl-UTP-Atto680, 1 μg BSA, and 2 μl (100 U) T7 RNA polymerase were incubated in 20 μl 1X RNA polymerase buffer for 1 hour at 37°C. 2 μl Turbo DNase (Life Technologies) and 2ul 10X Turbo DNase Buffer were added to the reaction and incubated at 37°C for 15 min. The RNA was denatured and separated on a 6% urea-polyacrylamide gel, cast on a Hoeffer miniVE apparatus and pre-run 20 minutes before loading. Full-length probe was excised, eluted overnight at 4°C in 0.5M Ammonium Acetate/1mM EDTA and ethanol precipitated. The concentration of the NIR labeled RNA probe was measured by absorption at 260 nm using the NanoDrop 1000 (Thermo Scientific).

The RNA competitors were generated by in vitro transcription. *pGEM-T Easy* was linearized with BsrBI to generate a 209 bp RNA competitor. *pGEM-Evf2(UCR)* was linearized with SalI to generate a 206 bp RNA competitor. The linearized templates were treated with Proteinase K and ethanol-precipitated. RNA was transcribed as follows: 1.25 μg DNA template, 10 mM DTT, 1.5 μl (80 U) RNasin (Promega), 2 mM A, C, G, and UTP (Roche), and 2 μl (100 U) T7 RNA polymerase (NEB) in 50 μl 1X RNA polymerase buffer were incubated at 37°C for 1 hour. Samples were incubated with 2μl Turbo DNase (Life Technologies) in 1X Turbo DNase Buffer for 15 minutes at 37°C. Samples were treated with Proteinase K (Roche), ethanol-precipitated and quantified using the Quantifluor RNA System (Promega).

To generate GST-tagged Sox2 proteins, full length Sox2 and Sox2 truncations (Sox2^1-205^, Sox2^1-109^, Sox2^206-317^, Sox2^41-317^) were subcloned into pGEX4T1. GST fusion proteins were purified from bacteria using standard protocols. The recombinant proteins were incubated with 0.15 pmoles *Evf2* NIR-labeled probe in 10 μl reactions for 30 min at room temperature. For all competition experiments, protein and competitor RNA were pre-incubated for 10 min at room temperature before adding probe. 5 μg tRNA and 0.5 μl RNasin (Promega) were included in all the REMSA reactions. Pre-electrophoresis of 4% native polyacrylamide gels was performed for 20 min, REMSA reactions loaded and electrophoresed at 200 V for 40 min, and data visualized in the Odyssey Infrared Imager (LI-COR Biosciences).

#### Primary embryonic brain MGE transfections

For all transfections, MGE tissues were dissected from E13.5 Swiss Webster embryos, dissociated in L15 media by pipetting several times, and spun through a cell strainer for single cell preparations. Cells were seeded at a density of 2.5 x 10^5^ cells per cm^2^ (Flandin et al. 2011) in neurobasal medium (DMEM/F-12 supplemented with L-glutamate, B-27 (Gibco), N2 supplement (Gibco), bovine pituitary extract (35 µg/mL; Life Technologies), mito+ serum extender (BD Biosciences), penicillin (100 U/mL; Gibco), streptomycin (100 µg/mL; Gibco), and glutamax (0.8 mM; Gibco)). One day prior to seeding cells, plate wells were coated with poly-L-lysine (Sigma) and laminin (Sigma).

For luciferase experiments, 78,300 cells per well were cultured in a 96-well microplate treated for tissue culture. Cells were allowed to attach for 24 hours before changing the medium to neurobasal media without antibiotics. Transfections using Fugene 6 were performed as recommended. Cells were harvested 48 hours after transfection with 1X passive lysis buffer (Promega) supplemented with 0.1% Digitonin for cell lysis. To ensure thorough cell lysis, lysates were subjected to two freeze-thaw cycles prior to performing Dual Luciferase Reporter assays. All transfections were normalized to the internal control expressing *Renilla luciferase,* performed at least in triplicate and a minimum of two times.

For transfection of mcherry-Sox2 fusions, 850,000 cells per well were cultured in a 12 well tissue culture plate. mcherry-Sox2 plasmids were generated by subcloning Sox2 or Sox2 truncations (Sox2^40-206^, Sox2^40-120^) into the mcherry2-C1 plasmid using SacI and KpnI. Quick Change Site Directed Mutagenesis (Agilent) mutagenesis was used to generate Sox2^Δ68-97^. 1 μg of plasmid was transfected using Fugene 6, as recommended. Cell were harvested by scrapping after 72 hr of incubation and nuclei isolated for combined RNA/DNA FISH and immunofluorescence using an anti-mcherry antibody (see method below).

#### Combined DNA-RNA FISH with Immunofluorescence

DNA FISH probes were generated by nick translation using the FISH Tag DNA Kit following manufacturer’s recommendations. The templates for the nick translation reactions were obtained from the BACPAC Resources Center (Children’s Hospital Oakland Research Institute): Dlx5/6 region: WI1-1693G2, Akr1b8 region: RP23-120B14, Rbm28 region: RP23-276H18. The digoxigenin labeled RNA probe was generated as described previously (Feng et al. 2006).

E13.5 whole ganglionic eminences were dissected in L15. Tissues were pooled for each genotype, triturated by pipetting, and filtered through a cell-strainer capped 5 ml polystyrene round-bottom tube (BD Falcon) to make single-cell suspensions. Cells were pelleted by centrifugation at 1000 rpm for 5 min at 4°C. The supernatant was removed and cells were gently resuspended in 500 μl Nuclear Extraction Buffer (0.32 M sucrose, 5 mM CaCl_2_, 3 mM Mg(Ac)_2_, 0.1 mM EDTA, 20 mM Tris-HCl pH 8.0, 0.1 % TritonX-100) and incubated on ice for 10 min. Cells were centrifuged at 100xg for 2.5 min at 4°C and the supernatant was removed. Cells were washed gently with ice-cold 1X PBS with 2 mM EGTA. Cells were centrifuged at 100 g for 2.5 min at 4°C. The supernatant was removed and cells were gently resuspended in 500 μl of ice-cold fixative (3 Methanol: 1 Glacial Acetic Acid). The cells were fixed for 10 min on ice. 5 μl of cells in fixative were transferred to Superfrost Plus microscope slides (Fisher Scientific) and allowed to air dry. The slides were transferred to a slide holder, vacuum-sealed and stored at -80°C.

Slides were incubated with 50 μg/mL pepsin in 0.01 M HCl at 37°C for 7 min, and washed twice with 2X SSC. Cells were fixed in 4% paraformaldehyde for 5 min at room temperature and washed 3 times with 2X SSC for 5 min. The slides were incubated in 1X PBS with 1% hydrogen peroxide for 30 min at room temperature and rinsed twice with 2X SSC. The slides were dehydrated by incubation for 2 min in 70%, 80% and 100% ethanol. 200 μl denaturation solution (70% formamide in 2X SSC) were added and the slides were incubated at 85°C for 10 min. Slides were dehydrated in ice-cold 70%, 80% and 100% ethanol for 2 min and allowed to air dry. 150 μl pre-hybridization buffer (50% formamide, 0.1% SDS, 300 ng/ml Salmon Sperm DNA, 2x SSC) were added and the slides were incubated overnight at 37°C.

DNA probes and RNA probe in hybridization buffer (50% formamide, 10% dextran sulfate, 0.1% SDS, 300 ng/ml Salmon Sperm DNA, 2x SSC) were denatured in the presence of 2 μg Mouse Hybloc DNA (Applied Genetics Laboratories) at 80°C for 7 min and re-annealed at 37°C for 1 hour. Slides were incubated for 5 min in 2X SSC with 50% formamide, 2 min in 4X SSC with 0.1% Tween-20 and 2 min in 2X SSC at 45°C. The slides were dehydrated in ethanol and denatured as described above. 10μl of FISH probe solution was added, and coverslips were sealed with rubber cement and the slides were incubated overnight at 37°C.

Slides were incubated in 2X SSC with 50% formamide for 10 min (3 times), in 2X SSC for 10 min and in 2X SSC with 0.1% NP40 for 5 min at 45°C. The slides were rinsed with 1X PBS and incubated in 1% blocking solution (Tyramide Signal Amplification Kit) for 1 hour. The appropriate antibody was diluted 1:500 in blocking reagent, along with a mouse monoclonal anti-Digoxigenin (DIG, 1:500), added to the slides and incubated at 4°C overnight. Slides were washed 3 times in 1X PBS for 3 min at room temperature, incubated with 1:100 HRP-goat anti-mouse IgG in blocking solution for 1 hour at room temperature and tyramide labeled according to manufacturer’s instructions (TSA Kit). A second tyramide labeling step was performed for immunostaining. Slides were washed 3 times in 1X PBS for 3 min at room temperature after the first round of labeling. Then the slides were incubated with 1:100 HRP-goat anti-rabbit IgG for 1 hour at room temperature and tyramide labeled. The slides were washed 3 times with 1X PBS for 3 min and incubated with 5 mg/ml DAPI for 5 min, rinsed with 1X PBS and mounted using *SlowFade* Gold antifade reagent (Thermo Fisher Scientific).

Cells were visualized using a Zeiss Laser Scanning Microscope 880 and a 100X immersion oil objective. Z-stacks of 0.3 μm intervals were obtained using the Zen 2.1 software. Imaris software was used for 3-D reconstruction, co-localization analysis and size measurements.

#### Cut& Run

Cut&Run was performed as previously described (Skene et al. 2018) with some modifications. The number of samples was determined according to the number of antibodies to be tested and the number of replicates (2 replicates per antibody and 2 replicates for IgG). The appropriate volume of Concanavalin A magnetic beads (10 μl per sample) were mixed into 1.5 ml of ice-cold Binding Buffer (20 mM HEPES-KOH pH 7.9, 10 mM KCl, 1mM CaCl_2_, 1mM MnCl_2_) and placed on a magnetic stand for 2 min. The supernatant was removed, and the beads were washed with 1.5 ml Binding Buffer. After magnetic separation of the beads, the supernatant was removed, and a volume of Binding Buffer equal to the initial beads volume (10 μl per sample) was added to the beads. The beads were placed on ice.

E13.5 ganglionic eminences were isolated from embryos in L15 medium (6 Embryos per genotype). Tissues were pooled for each genotype, triturated by pipetting, and filtered through a cell-strainer capped 5 ml polystyrene round-bottom tube (BD Falcon) to generate single-cell suspensions. Cells were counted using the Luna Automated Cell Counter (Logos Biosystems). The appropriate volume of cells to obtain 250,000 cells per sample was centrifuged at 600 g for 3 min at 4°C. The supernatant was removed and the cell pellet was gently resuspended in ice-cold Wash Buffer (20mM HEPES pH 7.5, 150 mM NaCl, 0.5 mM spermidine, EDTA-free protease inhibitor cocktail). The cells were centrifuged at 600 g for 3 min at 4°C. The supernatant was removed and cells were gently resuspended in 1 ml of Wash Buffer. The Concanavalin A bead suspension was added to the cells, while gently vortexing (∼1000 rpm), and the tube was incubated with rotation for 10 min at 4°C. The cells/beads suspension was split into aliquots, according to the number of samples previously determined. Tubes were placed on a magnetic stand and the supernatant removed. 50 ul of Antibody Buffer (20mM HEPES pH 7.5, 150 mM NaCl, 0.5 mM spermidine, 2 mM EDTA, 0.02% digitonin, EDTA-free protease inhibitor cocktail) containing 0.5 μg of antibody was added to each tube, while gently vortexing. The samples were incubated with rotation for 2 hours at 4°C. The samples were centrifuged for 5 seconds at 1000 rpm, and placed on the magnetic stand. The supernatant was removed, and the pellet resuspended gently in 1 ml ice-cold Digitonin Buffer (20 mM HEPES pH 7.5, 150 mM NaCl, 0.5 mM spermidine, 0.02% digitonin, EDTA-free protease inhibitor cocktail). The Digitonin Buffer wash was repeated once. Tubes were placed on the magnetic stand, the supernatant removed and 50 μl of pA-Mnase solution (final concentration 700 ng/ml in Digitonin Buffer) were added to each tube while gently vortexing. The samples were incubated with rotation for 1 hours at 4°C. The samples were centrifuged for 5 sec at 1000 rpm and placed on the magnetic stand. The supernatant was removed and the pellets were washed in ice-cold Digitonin Buffer two times, as described above. After removal of supernatant from washes, 150 μl of Digitonin Buffer were added to each sample while gently vortexing. Tubes were placed on a metal block on ice (0°C) for 5 min. Tubes were removed from ice briefly to add 3 μl 100 mM CaCl_2_ while gently vortexing, then tubes were returned to 0°C for 30 min. 100 ul of Stop Buffer (340 mM NaCl, 20 mM EDTA, 4mM EGTA, 0.02 % digitonin, 0.05 mg/ml RNAseA, 0.05 mg/m, 2 pg/ml heterologous spike-in DNA) were added to each sample, and mixed with gentle vortexing. Samples were incubated for 10 min at 37°C, then centrifuged for 5 min at 4°C at 16,000 rpm. The tubes were placed on the magnetic stand and the supernatant was transferred to a clean 1.5 ml microcentrifuge tube. DNA extraction was performed utilizing standard phenol chloroform and ethanol precipitation methods as described (Skene et al. 2018). Samples were ethanol precipitated overnight at -20°C. DNA pellets were dissolved in 20 ul 0.1x TE Buffer (1 mM Tris-HCl pH 8, 0.1 mM EDTA). The Qubit High-Sensitivity Assay was used for DNA quantification.

Cut&Run libraries were prepared using the KAPA Hyper Prep Kit protocol, with some modifications. The total volume of Cut&Run DNA was used for library construction. For adapter ligation, 5 μl of 3 μM Adapter stock from the KAPA Dual-Indexed Adapter Kit was used. The ligation was incubated at 20°C for 15 min. The library was amplified using the following cycling conditions: 98°C - 45 sec, 98°C -15 sec, 60°C -10 sec, 14 cycles, 72°C for 1 min. After library amplification, the libraries were purified using 50 ul of KAPA Pure Beads and eluted in 20 μl of water. Cut&Run sample quality was analyzed by TapeStation prior to sequencing on a NovaSeq 6000 (SP 100 cycles).

#### Cut&Run data processing

Paired-end Cut&Run reads of different transcription factors and their respective Ig-controls from Evf2+/+ and Evf2TS/TS Dlx5/6UCE samples were first mapped on mm9 genome using Bowtie2 v2.1.0 (Langmead and Salzberg 2012) with options ‘--local --very-sensitive-local --no-unal --no-mixed --no-discordant --phred33 -I 10 -X 700’. We used the Picard toolkit command ‘MarkDuplicates’ to mark PCR duplicates and remove them from the final mm9 genome mapped bam files. Next, we separated the sequence fragments into ≤ 120 and ≥ 150-bp size classes that provided the mapping of the local vicinity of a DNA-binding protein. The base-pair sizes can vary depending on the steric access to the DNA by the tethered MNase (Skene et al. 2018). Fragments mapping to repeat elements were removed, and replicates were joined before peak calling. The peak calling was performed using MACS2 (Zhang et al. 2008) callpeak options ‘-t -c –f BED -g mm --keep-dup all --bdg –nomodel --slocal 500 --llocal 5000 –-extsize 120/150’. An FDR cutoff of 0.05 was used to call the final set of peaks (Janssens et al. 2018). Differential Cut&Run analysis of transcription factor binding peaks between two Dlx5/6UCE conditions was performed using MACS2 program by treating one of the samples as the “control” for the other.

#### 4C-Seq

4C was performed as previously described (van de Werken et al. 2012), with some modifications. GEs from Sox2 ^fl/fl^ Cre+ and Sox2 ^fl/fl^ Cre-single embryos were dissected in L15 and kept in separate tubes. A single cell suspension was obtained through gentle pipetting of the tissue in 250 μl of L15. Cells (approximately 2 x10^6^ cells per embryo) were transferred to a tube containing 5 ml 2% paraformaldehyde/10% Fetal Bovine Serum (FBS) and incubated with rotation for 10 min at room temperature. 710 μl of 1M glycine were added to quench the formaldehyde and tubes were placed on ice. Cells were pelleted by centrifugation at 1300 rpm for 8 min at 4°C. The supernatant was removed and cells were gently resuspended in 2.5 ml ice-cold 4°C Lysis Buffer (50 mM Tris pH 7.5, 150 mM NaCl, 5 mM EDTA, 0.5% NP-40, 1% Triton X-100, protease inhibitors) and incubated for 10 min on ice. Cells were pelleted by centrifugation at 1800 rpm for 5 min at 4°C. The supernatant was removed, and the cells were washed with 1 ml ice-cold 1x PBS by gentle resuspension. Cells were transferred to 1.5 ml microcentrifuge tubes and centrifuged at 2,400 rpm for 2 min at 4°C. The supernatant was removed and the cell pellets were flash frozen in liquid nitrogen and stored at -80°C.

The cell pellets were resuspended in 440 μl of molecular grade water and 60 μl of Cutsmart Buffer (New England Biolabs) were added. Tubes were incubated at 37°C and 15 μl of 10% SDS were added. Samples were incubated for 1 hour at 37°C while shaking at 900 rpm. 75 ul of 20% Triton X-100 were added to the samples and incubated at 1 hour at 37°C while shaking at 900 rpm. 200 U of EcoRI-HF were added to the samples and incubated overnight at 37°C while shaking. The next day, 200 U of EcoRI-HF were added and incubated overnight at 37°C with shaking. Complete digestion was confirmed by agarose gel electrophoresis. If undigested DNA was still present, another 200 U of restriction enzyme was added and incubated overnight. The enzyme was inactivated at 65°C for 20 min. Samples were transferred to a 15 ml conical tube. 100 U of T4 DNA Ligase were added (2 ml reaction volume) and incubated overnight at 16°C. The next day, 500 μl of molecular grade water, 50 μl fresh T4 ligase buffer and 100 U of T4 DNA ligase were added to the samples and incubated overnight at 16°C. Complete ligation was confirmed by agarose gel electrophoresis. The DNA was extracted using standard ethanol precipitation procedures (van de Werken et al. 2012).

The DNA pellet was resuspended in 450 μl of molecular grade water, then 50 μl of DpnII buffer and 50 U of DpnII were added and incubated at 37°C overnight. Complete digestion was confirmed by agarose gel electrophoresis. The enzyme was inactivated at 65°C for 20 min. Samples were transferred to a 50 ml conical tube and ligation was performed using 100 U of T4 DNA ligase in a total volume of 3.5 ml at 16°C overnight. The DNA was precipitated using standard ethanol precipitation procedures. The samples were purified using the Qiaquick PCR purification kit (2 columns per sample).

The following steps were performed to generate the 4C library for sequencing. First, overhangs were added to the 4C template using PCR amplification with primers containing the bait sequence, as follows, PCR reaction: 200 ng 4C template, 0.2 mM dNTPs, 35 pmol Primer *Dlx5/6UCE*-Fwd, 35 pmol Primer *Dlx5/6UCE-*Rev, 1.75 U Expand Long Template Enzyme Mix (Roche), 1X Buffer I. PCR cycles: 94°C - 2 min, 94°C – 10 sec, 55°C - 1min, 68°C - 3 min, 29 cycles, 68°C - 5min. The PCR product was purified using the High Pure PCR Product Purification Kit (Roche). Then, the 4C DNA containing the overhangs was used as template for a second PCR that adds index sequences and Illumina sequencing adapters to generate the 4C library for sequencing. PCR reaction (50 μl): 225 ng DNA template, 0.5 mM dNTPs, 5μl Nextera XT Index1 primer (N7XX, Illumina), 5 μl Nextera Index 2 primer (S5XX, Illumina), 3.5 U Expand Long Template Enzyme Mix (Roche), 1X Buffer I. PCR cycles: 94°C - 5 min, 94°C – 10 sec, 55°C – 30 sec, 68°C - 1 min, 8 cycles, 68°C - 7min.The PCR product was purified using the High Pure PCR Product Purification Kit (Roche).

#### Dlx5/6UCE 4C-seq differential data analysis

4C reads were first mapped at the EcoRI restriction enzyme cut sites on chromosome 6 of mm9 reference genome using Bowtie2 v2.1.0 (Langmead and Salzberg 2012). The mapped reads were further filtered based on their reproducibility between the pair of replicates. An EcoRI cut-site was deemed to reproducibly interact (not interact) with the 4C bait if the two replicates in a given condition (*Evf2^+/+^* and *Evf2^TS/TS^*) both have non-zero (zero) counts. We identified 1108 and 1266 non-zero count EcoRI restriction cut-sites that are reproducible in both replicates of *Evf2^+/+^* and *Evf2^TS/TS^*, respectively. Across the two conditions (*Evf2^+/+^* and *Evf2^TS/TS^*), we retained a total of 997 reproducible 4C sites that has reproducible interactions in the two replicates of either one condition or in both conditions. We then performed a DESeq2 (Love et al. 2014) based differential contact count analysis on these sites to identify Evf2-regulated sites (p-adjusted value < 0.05 and a log2 fold change ≥ 2 for positively regulated (**+**) or ≤ -2 for negatively regulated (**-**)) and Evf2-independent (**I**) (p-adjusted value > 0.05 and an absolute log2 fold change < 2) 4C interaction sites. We also performed the 4Cseq analysis using the FourCSeq program (Klein et al. 2015). FourCSeq program models the overall decreasing interaction frequency with genomic distance by fitting a smooth monotonically decreasing function to suitably transformed count data. With this transformed and normalized count data, FourCSeq performs differential analysis between conditions to get significant differential interactions. We applied FourCSeq on our *Evf2^+/+^* and *Evf2^TS/TS^* 4Cseq samples and retrieved (Dlx5/6UCE+) and (Dlx5/6UCE-) Evf2 regulated 4C interactions (p-adjusted value < 0.05 and an absolute log2 fold change ≥ 2) and Dlx5/6UCE-I, Evf2-independent interactions (p-adjusted value > 0.05 and an absolute log2 fold change < 2). To avoid method-specific biases, interaction sites that were assigned the same label (+, -, I) by the two different approaches (DESeq2 and FourCSeq) were called 4Cseq-intersectional computational method sites (4Cseq-ICMs).

#### Sox2flfl +/cre 4C-seq differential data analysis

4C reads were first mapped at the EcoRI restriction enzyme cut sites on chromosome 6 of mm9 reference genome using Bowtie2 v2.1.0 (Langmead and Salzberg 2012). 4C contact counts were generated from the Bowtie2 mapped bam files using FourCSeq program (Klein et al. 2015). We then performed a DESeq2 (Love et al. 2014) based differential contact count analysis on these sites to identify Sox2flfl sites (p-adjusted value < 0.05 and a log2 fold change ≥ 2 for positively regulated (cre) or ≤ -2 for negatively regulated (+) 4C interaction sites with non-zero counts in at least two of the replicates in a differential condition. We also performed the 4Cseq analysis using the FourCSeq program. FourCSeq program models the overall decreasing interaction frequency with genomic distance by fitting a smooth monotonically decreasing function to suitably transformed count data. With this transformed and normalized count data, FourCSeq performs differential analysis between conditions to get significant differential interactions. We applied FourCSeq on our *Sox2flfl +* and *Sox2flfl cre* 4Cseq samples and retrieved Sox2 regulated 4C interactions (p-adjusted value < 0.05 and an absolute log2 fold change ≥ 2). To avoid method-specific biases, we retained a common set of 244 4C differential sites.

#### Dlx5/6UCE 4C-seq and Cut&Run signal overlap

*Evf2^+/+^* and *Evf2^TS/TS^ Dlx5/6UCE* 4C-seq peaks were first mapped on Cut&Run differential transcription factor peaks using bedtools ‘intersect’ function (Quinlan and Hall 2010). The overlapping set of 4C-seq peaks from a *Dlx5/6UCE* condition were then mapped on the respective Cut&Run Ig-normalized transcription factor signal data. The log2 fold-enrichment of Ig-normalized signal was generated using MACS2 ‘bdgcmp’ command (Zhang et al. 2008). The violin plots were made using ‘ggpubr’ R package (Kassambara).

#### Chromatin immunoprecipitation (ChIP)

For ChIP experiments whole ganglionic eminences were dissected from 10 embryos per genotype. Tissues were pooled for each genotype, triturated by pipetting, and filtered through a cell-strainer capped 5 ml polystyrene round-bottom tube (BD Falcon) to make single-cell suspensions. Duplicate ChIP experiments were performed to determine reproducibility, generating libraries as described below.

#### Cross-linked ChIP

For anti-DLX and anti-H3K27ac ChIP cells were fixed in 1% paraformaldehyde for 10 min. For anti-Sox2, anti-Smc1 and anti-Smc3 ChIP cells were fixed in 1% paraformaldehyde for 90 min. Cells were lysed in SDS lysis buffer (1% SDS, 50 mM Tris-HCl pH 8, 10 mM EDTA) with protease inhibitors. The lysates were sonicated with a Bioruptor Pico (Diagenode) for 10 cycles (30 sec On, 30 sec Off). The lysates were then centrifuged to pellet cellular debris and the supernatant collected for ChIP. 25 μg of chromatin were diluted 1:10 in RIPA Buffer (10mM Tris pH 7.6, 1mM EDTA, 0.1% SDS, 0.1% Sodium Deoxycholate, 1% Triton X-100) with protease inhibitors. The chromatin was pre-cleared by rotating at 4°C with 50 μl of Protein G–Agarose beads for 1 hour. After centrifugation to pellet the beads, the supernatant was further pre-cleared by rotating at 4°C with 50 μl rabbit IgG conjugated Protein G–Agarose beads for 1 hour. The precleared chromatin was incubated with rabbit IgG (2.5 μg), previously validated anti-pan-DLX (2.5 μg), anti-H3K27ac (1 μg), anti-SMC1 (1 μg), anti-SMC3 (1 μg) or anti-Sox2 (μg) at 4°C for 4 hours with rotation. 50 μl of Protein G-Agarose beads blocked with 1% BSA in 1X PBS were added to each sample and incubated at 4°C overnight with rotation. Beads were pelleted by centrifugation and washed twice with Low Salt Buffer (20 mM Tris-HCl pH 8.1, 2 mM EDTA, 150 mM NaCl, 0.1% SDS, 1% Triton X-100), three times with High Salt Buffer (20 mM Tris-HCl pH 8.1, 2 mM EDTA, 500 mM NaCl, 0.1% SDS, 1% Triton X-100), four times with LiCl buffer (0.25M LiCl, 10 mM Tris-HCl pH 8.1, 1 mM EDTA, 1% sodium deoxycholate and 1% NP-40), twice with 0.1% Tween-20 in 1X PBS, and once with TE buffer (10 mM Tris-HCl pH 8.1 and 1 mM EDTA). Immunoprecipitated DNA was eluted from the beads by incubation with 200 μl of elution buffer (50 mM Tris-HCl pH 8, 10 mM EDTA, 1% SDS) at 65°C for 1 hour. The beads were removed by centrifugation and DNA crosslinking was reversed at 65°C for 4 hours. The DNA was incubated with 20 mg of RNAse A at 55°C for 1 hour. 40 mg Proteinase K were added and incubated at 55°C for 1 hour. The Immunoprecipitated DNA was purified using the Qiaquick PCR Purification Kit (Qiagen).

#### ChIP-re-ChIP

For ChIP-re-ChIP cells were fixed in 1% paraformaldehyde for 90 min, then lysed in SDS lysis buffer (1% SDS, 50 mM Tris-HCl pH 8, 10 mM EDTA) with protease inhibitors. The lysates were sonicated with a Microson Ultrasonic Cell Disruptor using 6 pulses of 10 seconds at 9 watts (RMS). The lysates were then centrifuged to pellet cellular debris and the supernatant collected for ChIP.

25 μg of chromatin were diluted 1:10 in RIPA Buffer (10mM Tris pH 7.6, 1mM EDTA, 0.1% SDS, 0.1% Sodium Deoxycholate, 1% Triton X-100) with protease inhibitors. The chromatin was pre-cleared by rotating at 4°C with 50 μl of Protein G–Agarose beads for 1 hour. After centrifugation to pellet the beads, the supernatant was further pre-cleared by rotating at 4°C with 50 μl rabbit IgG conjugated Protein G–Agarose beads for 1 hour. The pre-cleared chromatin was incubated with rabbit IgG (5 μg), anti-DLX (5 μg), at 4°C for 4 hours with rotation. 50 μl of Protein G-Agarose beads blocked with 1% BSA in 1X PBS were added to each sample and incubated at 4°C overnight with rotation. Beads were pelleted by centrifugation and washed 3 times with re-ChIP Buffer (20 mM EDTA

0.5M NaCl, 0.1% SDS, 500µl NP40) and two times with 1X TE buffer. Beads were pelleted by centrifugation and the supernatant removed. 75 ul re-ChIP elution buffer (2% SDS,

15mM DTT, protease inhibitors) were added to the beads and samples were incubated at 37°C for 30 min. Beads were pelleted by centrifugation and the supernatant transferred to a clean tube. ChIP elution buffer supplemented with 50 ug BSA was added to the samples to a total volume of 1.5 ml. 1 ug anti-Sox2 or anti-LaminB1 antibody were added to the samples and incubated for 2-4 hours at 4°C with rotation. 50 μl of BSA blocked Protein G-Agarose beads were added to each sample and incubated at 4°C overnight with rotation.

Beads were pelleted by centrifugation and washed once with each of the following buffers: Low Salt Buffer (20 mM Tris-HCl pH 8.1, 2 mM EDTA, 150 mM NaCl, 0.1% SDS, 1% Triton X-100), High Salt Buffer (20 mM Tris-HCl pH 8.1, 2 mM EDTA, 500 mM NaCl, 0.1% SDS, 1% Triton X-100), LiCl buffer (0.25M LiCl, 10 mM Tris-HCl pH 8.1, 1 mM EDTA, 1% sodium deoxycholate and 1% NP-40), 0.1% Tween-20 in 1X PBS, and TE buffer (10 mM Tris-HCl pH 8.1 and 1 mM EDTA). Immunoprecipitated DNA was eluted from the beads by incubation with 200 μl of elution buffer (50 mM Tris-HCl pH 8, 10 mM EDTA, 1% SDS) at 65°C for 1 hour. The beads were removed by centrifugation and DNA crosslinking was reversed at 65°C for 4 hours. The Immunoprecipitated DNA was purified using the Qiaquick PCR Purification Kit (Qiagen).

#### Native ChIP

Native ChIP was performed for the following antibodies: H3K4me1, H3K4me3, H3K27me3 as previously described (Cajigas et al. 2018). Cells from the single cell suspension described above were split into 1 × 10^6^ cell aliquots, and pelleted through centrifugation at 1000 × g for 10 min. Cell pellets were flash frozen in liquid nitrogen, and stored at −80°C. Nuclei were isolated using EZ Nuclei Isolation Lysis Buffer (Sigma). Chromatin was digested in 2U/μl Micrococcal nuclease (NEB) at 37°C for 7 min. The reaction was quenched with EDTA (10 mM final concentration). Triton X-100 and Sodium Deoxycholate were added (0.1% final concentration). Samples were incubated on ice for >15 minutes. Immunoprecipitation buffer (20 mM Tris-HCl pH 8.0, 2 mM EDTA, 150 mM NaCl, 0.1% Triton X-100, 1x Protease inhibitor cocktail, 1 mM PMSF) was added to a final volume of 200 μl and the samples were rotated at 4°C for 1 hour. The chromatin was pre-cleared by rotating at 4°C with 15 μl of Protein G–Agarose beads for 1 hour. After centrifugation to pellet the beads, the supernatant was further pre-cleared by rotating at 4°C with 15 μl rabbit IgG conjugated Protein G–Agarose beads for 1 hour. The pre-cleared chromatin was incubated with rabbit IgG (1 μg), or antibodies targeting histone modifications (1 μg) at 4°C for 1–2 hours with rotation. 15 μl of Protein G-Agarose beads blocked with 1% BSA in 1X PBS were added to each sample and incubated at 4°C overnight with rotation. The beads were pelleted by centrifugation and washed twice with 200 μl Low Salt Wash buffer (20 mM Tris-HCl pH 8.0, 2 mM EDTA, 150 mM NaCl, 1% Triton X-100, 0.1% SDS) and twice with 200 μl High Salt Wash buffer (20 mM Tris-HCl pH 8.0, 2 mM EDTA, 500 mM NaCl, 1% Triton X-100, 0.1% SDS). Immunoprecipitated DNA was eluted in 100 μl of ChIP elution buffer (100 mM NaHCO3, 1% SDS) at 65°C for 1–1.5 hour. The DNA was purified using phenol chloroform extraction and ethanol precipitated. The pellet was resuspended in 10 mM Tris-HCl pH 8.5. The DNA was incubated with 20 mg of RNAse A at 55°C for 1 hour. 40 mg Proteinase K were added and incubated at 55°C for 1 hour. The immunoprecipitated DNA was purified using the Qiaquick PCR Purification Kit.

The Quant-iT PicoGreen dsDNA Assay Kit was used for quantification of ChIP samples. 150ng to 1 μg of DNA was prepared into Illumina libraries, according to manufacturer’s instructions, using the TruSeq Nano DNA Library Prep Kit. Resulting libraries were deep sequenced, using the Illumina HiSeq2500 system in Rapid Run mode, obtaining between 10M and 15M of 100-bp length, single-end reads per library.

#### ChIP-seq read alignment

Raw sequencing reads for all the individual ChIP-seq datasets were aligned using bwa mapper (Li and Durbin 2009) version 0.7.12 with the following settings ‘aln -t 8 samse’. We allowed two mismatches relative to the reference and only retained the unique alignments with Phred quality score greater than 30. The datasets were mapped against mm9 version of the mouse genome.

#### ChIP-seq data analysis

##### Quality assessment

ChIP-seq quality assessment was carried out using the strategy described by ENCODE ChIP-seq data analysis guidelines (Landt et al. 2012). Cross-correlation analysis was performed using SPP package (Kharchenko et al. 2008) using the parameter ‘-s = - 100:5:600’. The analysis is essential to assess the NSC (Normalized Strand Correlation) and RSC (Relative Strand Correlation) values as recommended by ENCODE (Landt et al. 2012). As per the guideline, all of our selected ChIP-seq datasets are above NSC value (> 1.05) and RSC value (> 0.8) threshold, and subsequent QC scores equal to or above 1 (Landt et al. 2012).

##### Peak calling and differential ChIP-seq analysis

After quality assessment, we used ‘‘irreproducible discovery rate’’ (IDR) framework to call the peaks against their respective input ChIP libraries using MACS2 program (Zhang et al. 2008) as described in the ENCODE guidelines (Landt et al. 2012). MACS2 peak calling was performed using the following settings ‘-p 1e-3 –to-large –nomodel –shiftsize’ while the other parameters were set to their default mode. The final conservative set of peaks for all the samples were called across technical replicates with an IDR threshold of 0.01.

#### Differential ChIP-seq analysis

Differential ChIP-seq analysis between two conditions was performed using MACS2 program (Zhang et al. 2008) by treating one of the samples as the “control” for the other. The peak identification by MACS2 was carried out using the same parameter settings as previously described in the ChIP-seq data analysis section. The cross-correlation analysis step (Kharchenko et al. 2008) was also performed on the respective datasets to determine the ‘–shiftsize’ parameter essential for peak identification by MACS2.

### QUANTIFICATION AND STATISTICAL ANALYSIS

#### General data analysis

Quantification and statistical analysis were performed using R. Significance levels: ∗ < 0.05, ∗∗ < 0.01, ∗∗∗ < 0.001. Violin plot: horizontal black line depicts the median, the box is the 25^th^ and 75^th^ percentile and the whiskers denote the minimum and maximum values considered for analysis. The colored dotted line represents the mean of the respective class. An unpaired T-test was used to measure the significance. For ChIP-seq, and CUT&RUN a peak is defined as a region with q-value less than 0.05 while for a 4C-seq experiment a significant peak is defined with FDR < 0.05 and an absolute log2 fold enrichment of ≥ 2. Addition statistical details can be found in the figure legends.

## KEY RESOURCES TABLE

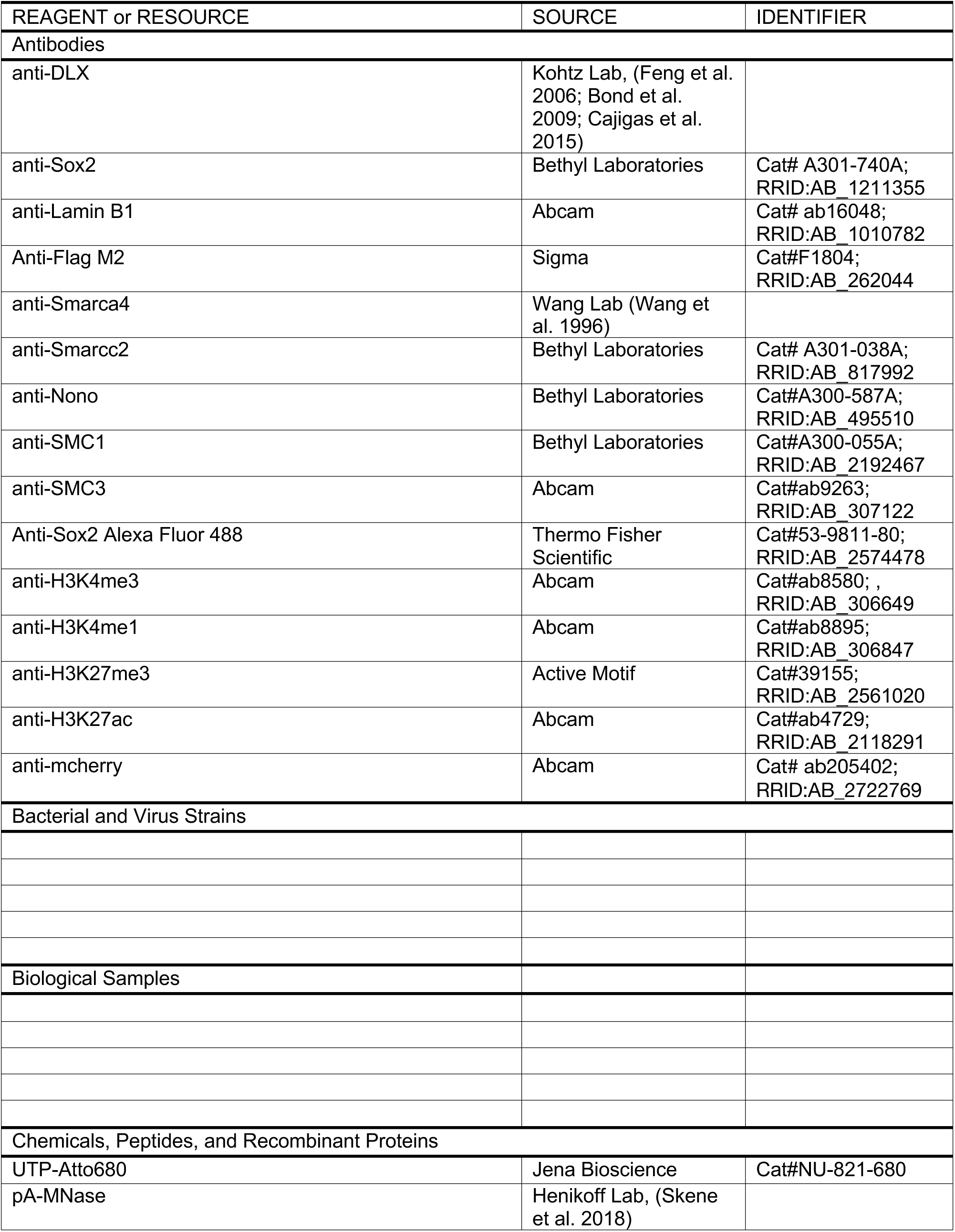
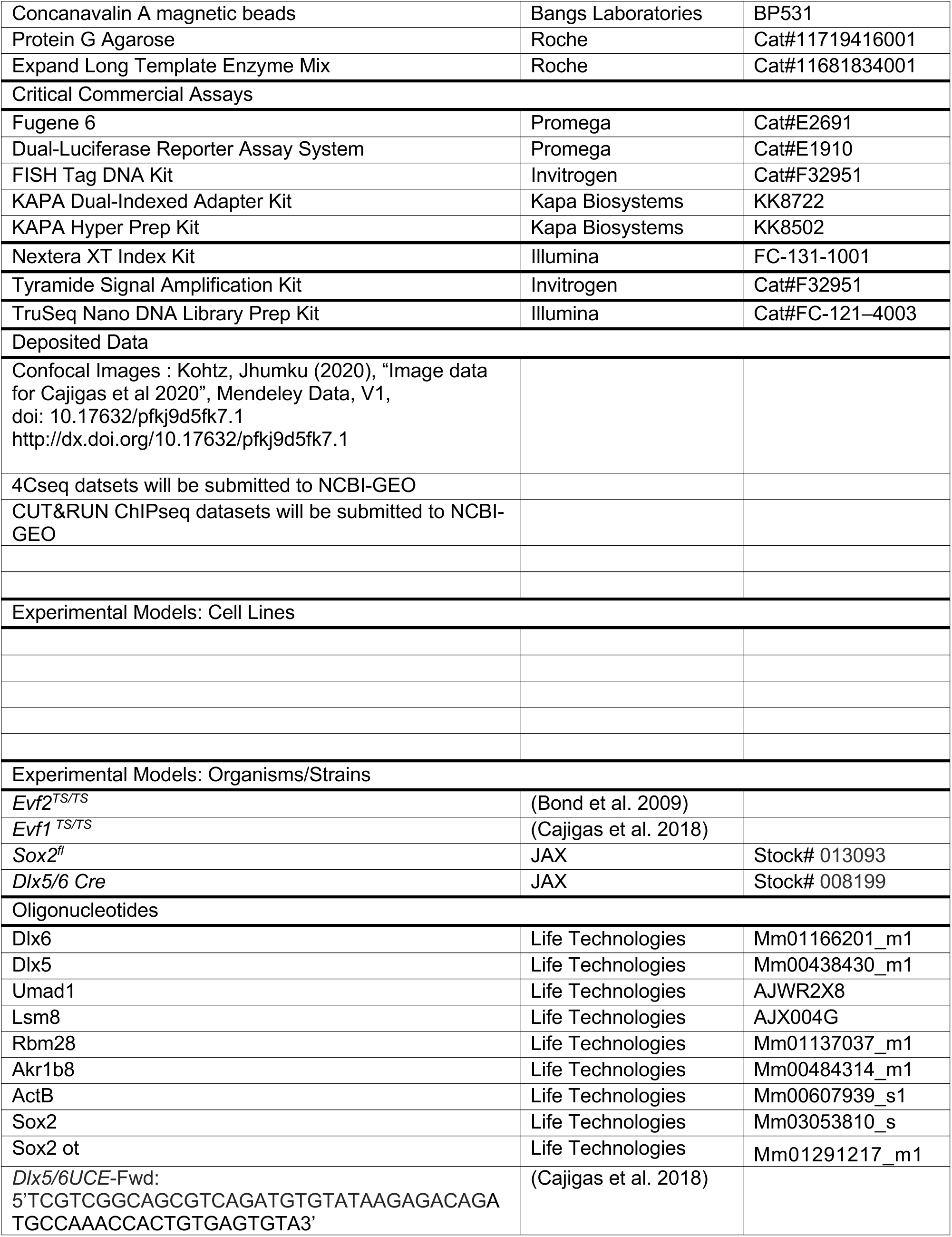
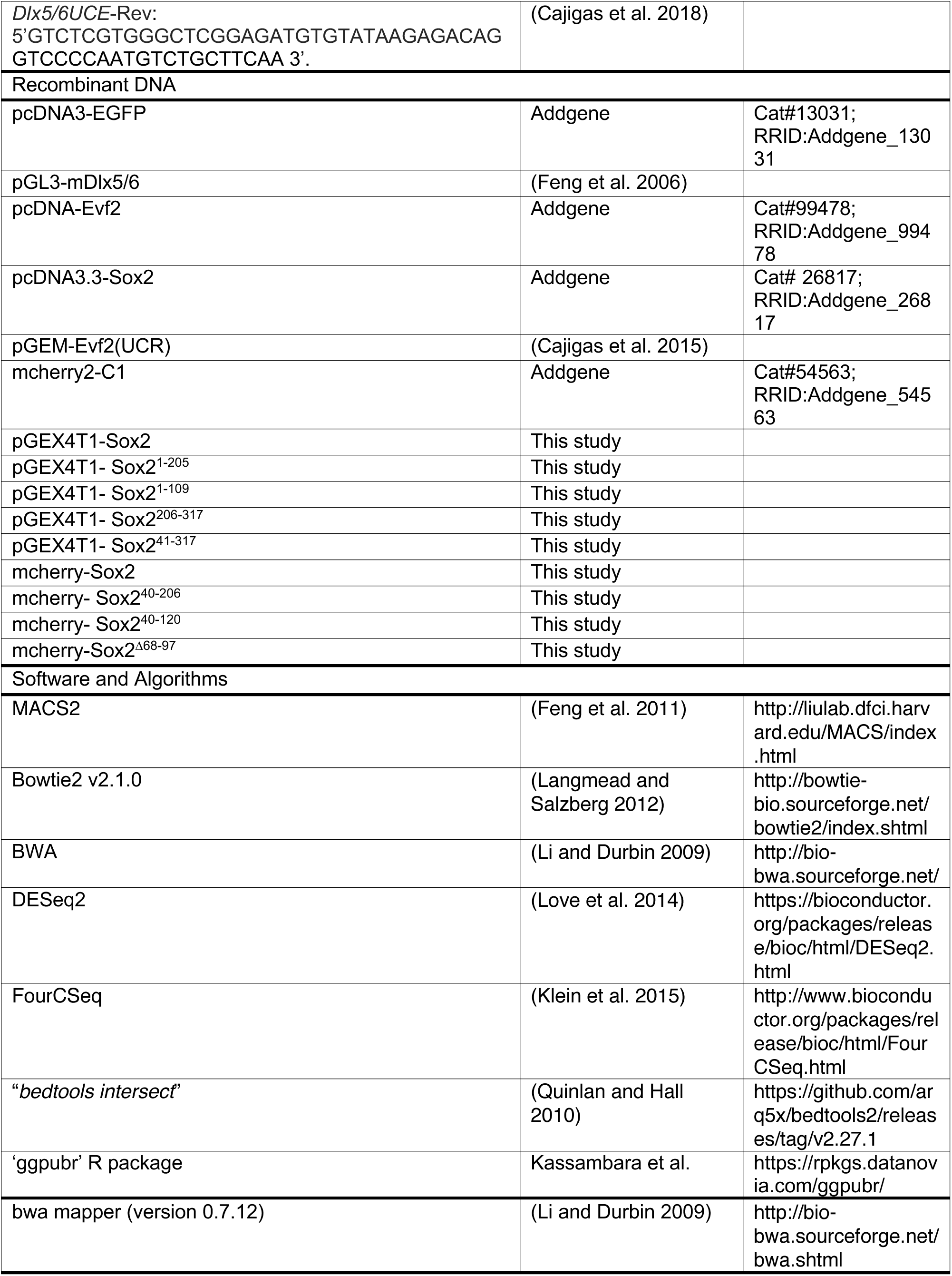
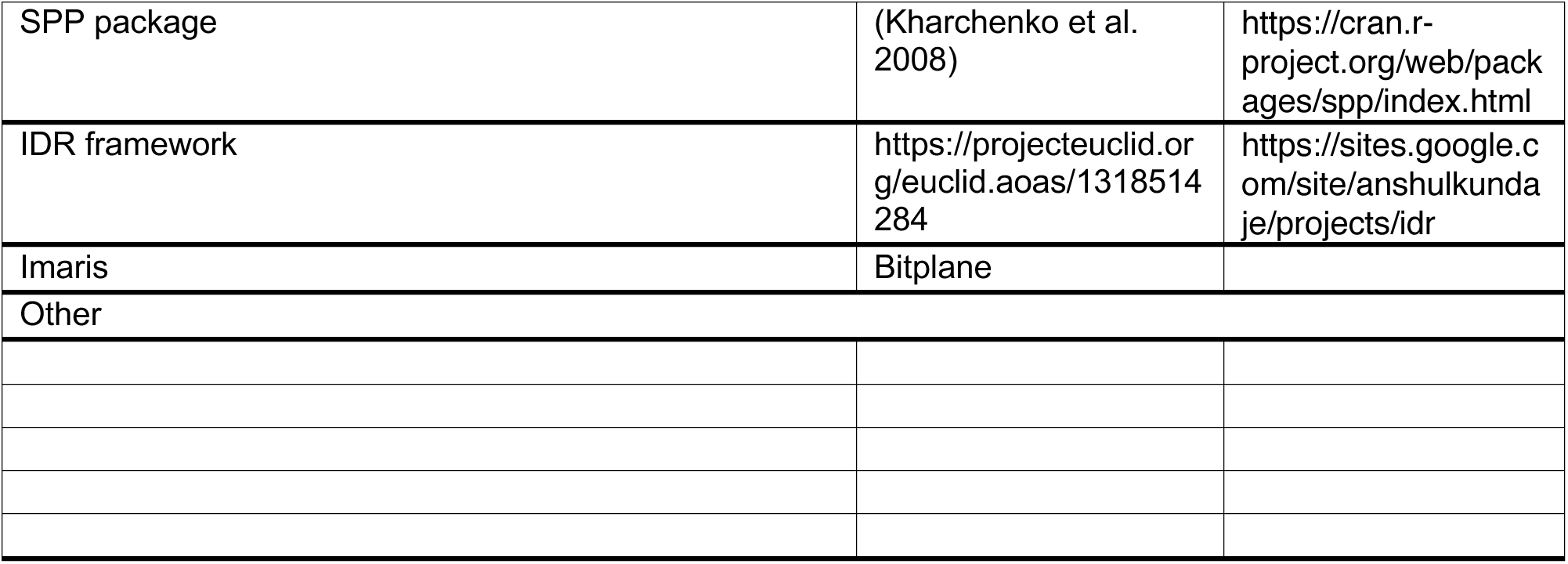

## Supplemental Figures

**Figure S1.**
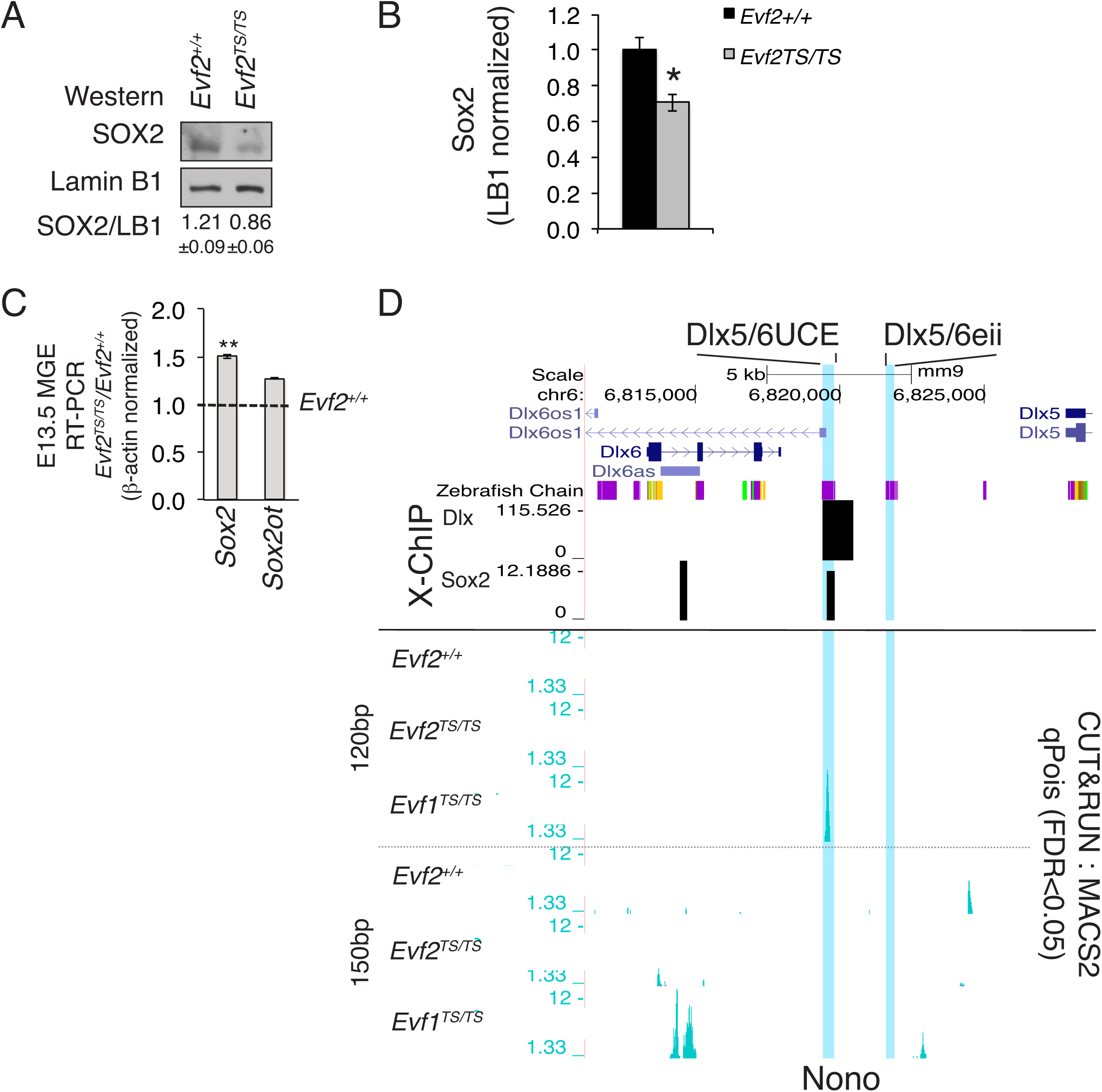
*Evf2* regulation of *Evf2* RNPs Sox2 and Nono. **A. B**. Western analysis of Sox2 protein levels in *Evf2* expressing (*Evf2^+/+^*) and *Evf2* lacking (*Evf2^TS/TS^*) E13.5GE extracts, (normalized to Lamin B1) shows ∼25% decrease in Sox2 protein levels in the absence of *Evf2*. **C.** Taqman qRT-PCR of Sox2 and Sox2ot RNA levels in *Evf2^+/+^* and *Evf2^TS/TS^* E13.5GE shows ∼50% increase in Sox2 RNA, but no change in Sox2ot RNA (normalized to β-actin). A-C. n=3/genotype, Student’s *t* test. **D.** UCSC browser display of CUT&RUN native ChIPseq binding profiles of *Evf2*-RNP Nono in *Evf2^+/+^*, *Evf2^TS/TS^*, *Evf1^TS/TS^* GE. *Dlx5/6* intergenic enhancers (*Dlx5/6UCE* and *Dlx5/6eii* shadow enhancer) are highlighted in blue. 120bp and 150bp fragment profiles show MACS2 validated peaks (FDR <0.05). to *Dlx5/6UCE,* distin-guishing Nono from other RNPs in these studies. Nono peak at *Dlx5/6UCE* is detected upon *Evf2* trun-cation (*Evf1^TS/TS^*), supporting that *Evf2-3’* inhibits Nono recruitment n=4/each genotype.

**Figure S2.**
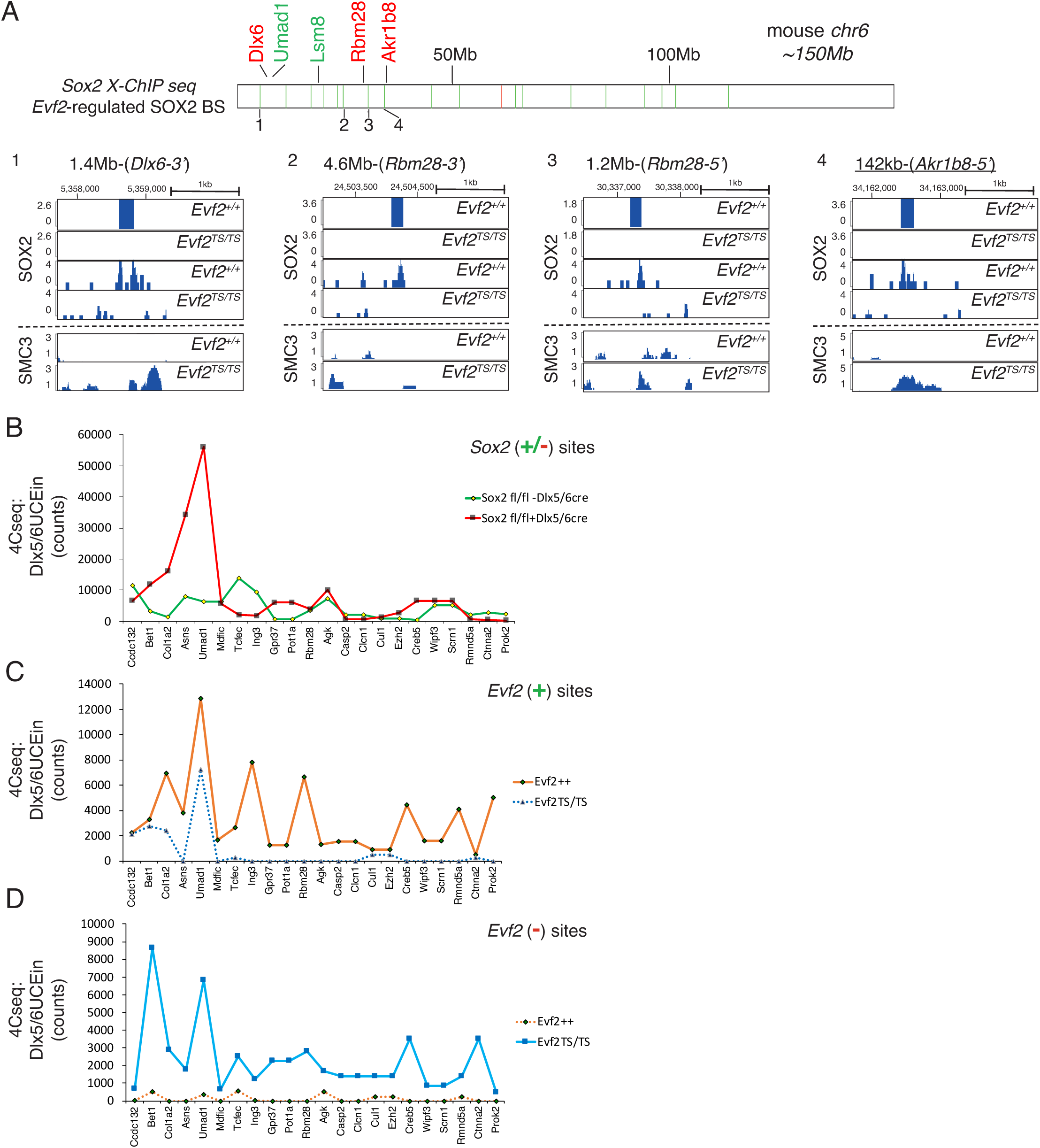
*Evf2* regulation of *Evf2* RNPs Sox2 and Smc3. **A.** Sox2 ChIPseq profiles across mouse chr6 from crosslinked chro-matin isolated from *Evf2* expressing (*Evf2^+/+^*) and *Evf2* lacking (*Evf2^TS/TS^*) E13.5GE. *Evf2* positively regulated Sox2 binding sites (BS) (green bars), and negatively regulated (red bars) are shown. *Evf2* repressed gene tar-gets are labeled in red (Dlx, Rbm28, Akr1b8), activated gene targets in green (Umad1, Lsm8). Peaks from sites labelled 1-4 overlap with Evf2 positively regulated Sox2 binding (differential MACS2 binding site shown in top two rows) and Smc3 bind-ing peaks. Site 1: Sox2 (+) Smc3 (-), Site 2: Sox2 (+) Smc3 (no peak), Site 3: Sox2 (+) Smc3 binding (but not Evf2 regulated), Site 4:Sox2 (+) Smc3 (-), n=2/genotype, Sites 1- 4 top two rows (differential MACS2), rows 3- 6 (MACS2. FDR<0.05). **B-D**. 4Cseq counts near genes across mouse chr6 identify over-lapping *Evf2* regulated and Sox2 regulated *Dlx5/6UCEins* from E13.5GEs **B**. *Sox2* positively regulated *Dlx5/6UCE*ins from Sox2fl/fl;-Dlx5/6cre (presence of Sox2), and Sox2fl/fl;+Dlx5/6cre (absence of Sox2). **C.** *Evf2* positively regulated *Dlx5/6UCEin* sites (orange) *Evf2^+/+^* (presence of *Evf2*), and *Evf2^TS/TS^* (absence of *Evf2*). **D**. *Evf2* nega-tively regulated sites (blue) *Evf2^+/+^* (pres-ence of *Evf2*), and *Evf2^TS/TS^* (absence of *Evf2*). B-D. n=4/each genotype 4Cseq inter-section of two computational methods (FourCseq and DEseq).

**Figure S3.**
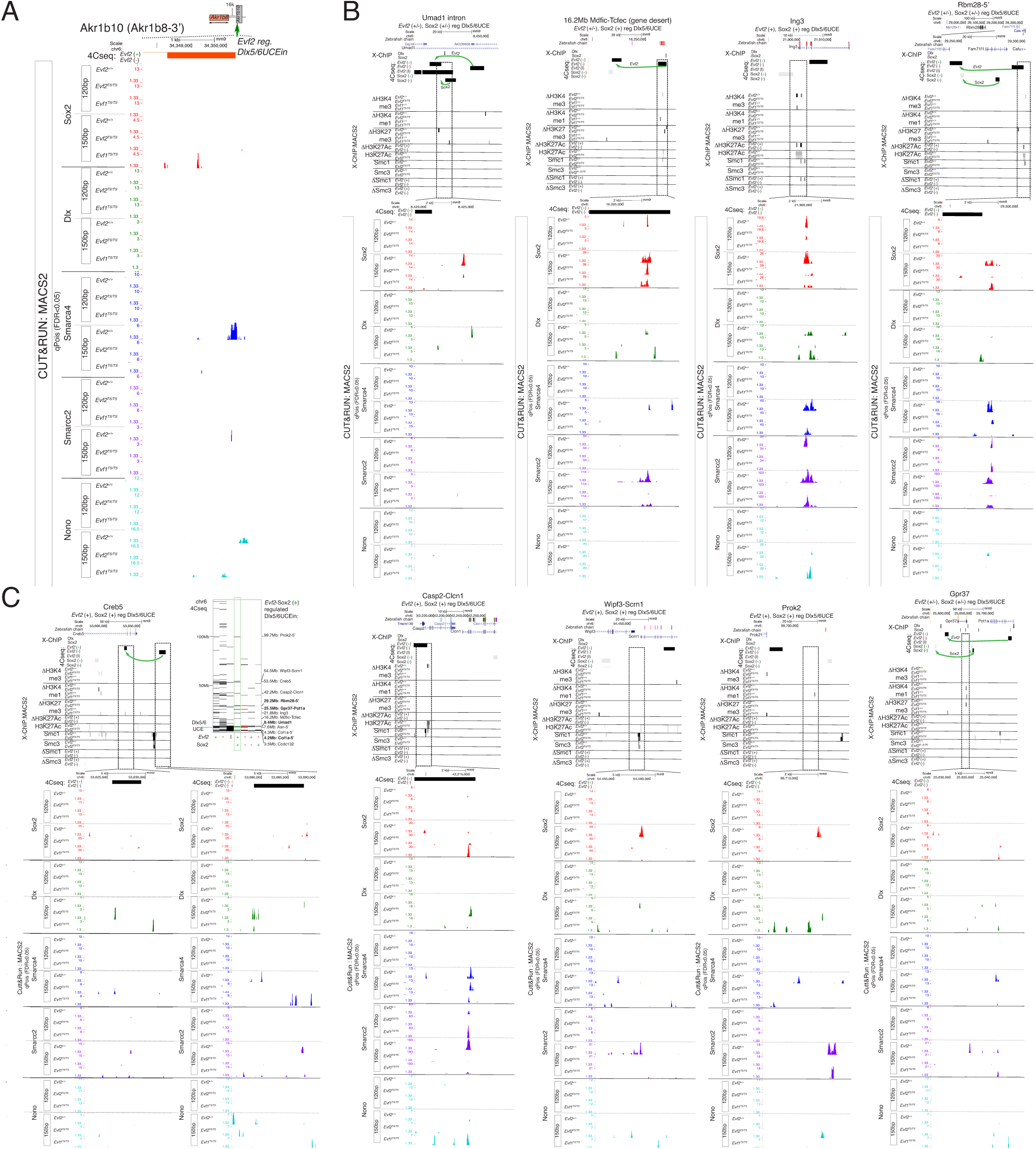
Relationships between *Evf2*-regulated *Dlx5/6UCEin*, RNP binding, and histone modifications at specific sites on mouse *chr6*. CUT &RUN profiles of *Evf2*-regulated binding *Evf2* RNPs Sox2, Dlx, Smarca4 and Smarcc2, and histone modifications (H3K27ac/me3, H3K4me3, H3K4me1) at select *Evf2*-regulated *Dlx5/6UCEin*s from *Evf2^+/+^*, *Evf2^TS/TS^*, and *Evf1^TS/TS^* E13.5GE. **A.** *Evf2* regulates Smarca4, but not Sox2 at the *Evf2* positively regulated *Dlx5/6UCEin* located 16kb 3’ of *Evf2* repressed target gene Akr1b8. **B-C.** ChIPseq profiles of histone modifications (crosslinked and native chromatin), Smc1a/Smc3 (crosslinked chromatin), Sox2, Dlx, Smarca4, Smarcc2, Nono (CUT&RUN) overlapping with Evf2 regulated Dlx5/6UCEins generated from 4Cseq analysis. ChIPseq MACS2 peak (FDR<0.05), n=2-4/genotype, 4Cseq FourCseq intersection with DEseq, n=4/genotype

**Figure S4.**
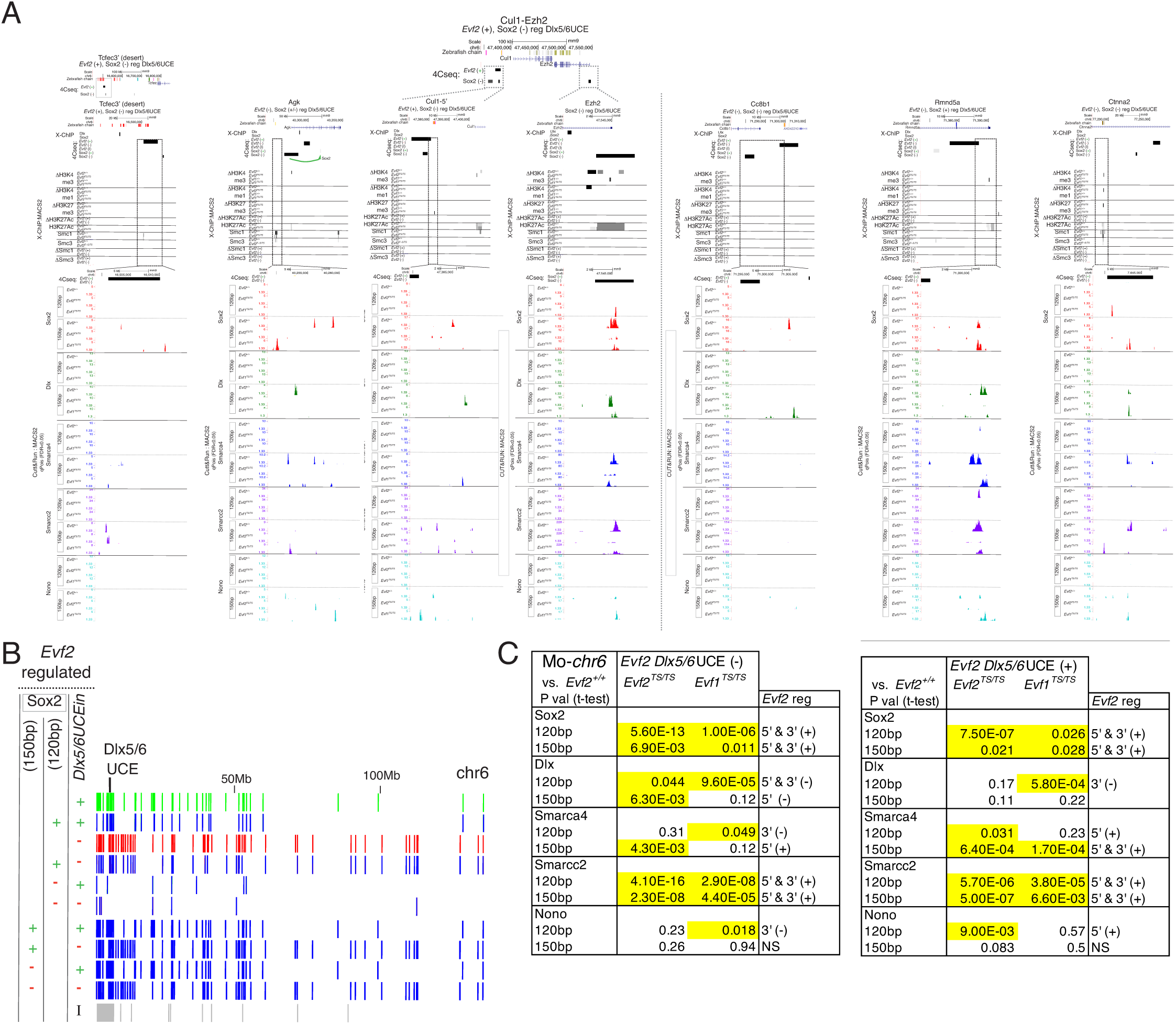
*Evf2*-regulated *Dlx5/6UCEin*, RNP binding, and histone modifications on mouse *chr6*. CUT &RUN profiles of *Evf2*-regulated *Evf2* RNPs Sox2, Dlx, Smarca4 and Smarcc2, histone modifications (H3K27ac/me3, H3K4me3, H3K4me1) and *Evf2*-regulated *Dlx5/6UCEin*s from *Evf2^+/+^*, *Evf2^TS/TS^*, and *Evf1^TS/TS^* E13.5GE. **A.** Relationships between ChIPseq profiles of histone modifications (crosslinked and native chromatin), Smc1a/Smc3 (crosslinked chromatin), and Sox2, Dlx, Smarca4, Smarcc2, Nono (CUT&RUN, native chromatin) overlapping with *Evf2* regulated *Dlx5/6UCEins* generated from 4Cseq analysis. **B.** *Evf2* regulated Sox2 binding sites overlapping with *Evf2* positively reg-ulated (+) and negatively regulated (-) and independent *Dlx5/6UCEins* across *chr6*. **C**. Statistically significant differences (yellow highlights) in RNP binding (CUT&RUN ChIPseq) at *Evf2* positively and negatively regulated *Dlx5/6UCEins* (4Cseq) from *Evf2^+/+^*, *Evf2^TS/TS^*, and *Evf1^TS/TS^* E13.5GE.

